# TBCK-deficiency leads to compartment-specific mRNA and lysosomal trafficking defects in patient-derived neurons

**DOI:** 10.1101/2025.03.02.641041

**Authors:** Marco Flores-Mendez, Jesus A. Tintos-Hernández, Leonardo Ramos-Rodriguez, Leann Miles, Tsz Y Lo, Yuanquan Song, Xilma R. Ortiz-González

**Affiliations:** Department of Pediatrics, Division of Neurology, The Children’s of Philadelphia, Philadelphia, PA; Center for Mitochondrial and Epigenomic Medicine, The Children’s Hospital of Philadelphia, Philadelphia, PA; Department of Biomedical Graduate Studies, Perelman School of Medicine, University of Pennsylvania, Philadelphia, PA; Raymond G. Perelman Center for Cellular and Molecular Therapeutics, The Children’s Hospital of Philadelphia, Philadelphia, PA; Department of Pathology and Laboratory Medicine, University of Pennsylvania, Philadelphia, PA; Department of Neurology, Perelman School of Medicine, University of Pennsylvania, Philadelphia, PA

## Abstract

Monogenic pediatric neurodegenerative disorders can reveal fundamental cellular mechanisms that underlie selective neuronal vulnerability. TBCK-Encephaloneuronopathy (TBCKE) is a rare autosomal recessive disorder caused by stop-gain variants in the *TBCK* gene. Clinically, patients show evidence of profound neurodevelopmental delays, but also symptoms of progressive encephalopathy and motor neuron disease. Yet, the physiological role of TBCK protein remains unclear. We report a human neuronal TBCKE model, derived from iPSCs homozygous for the Boricua variant (p.R126X). Using unbiased proteomic analyses of human neurons, we find TBCK interacts with PPP1R21, C12orf4, and Cryzl1, consistent with TBCK being part of the FERRY mRNA transport complex. Loss of TBCK leads to depletion of C12ORF4 protein levels across multiple cell types, suggesting TBCK may also play a role regulating at least some members of the FERRY complex. We find that TBCK preferentially, but not exclusively, localizes to the surface of endolysosomal vesicles and can colocalize with mRNA in lysosomes. Furthermore, TBCK-deficient neurons have reduced mRNA content in the axonal compartment relative to the soma. TBCK-deficient neurons show reduced levels of the lysosomal dynein/dynactin adapter protein JIP4, which functionally leads to TBCK-deficient neurons exhibiting striking lysosomal axonal retrograde trafficking defects. Hence, our work reveals that TBCK can mediate endolysosomal trafficking of mRNA, particularly along lysosomes in human axonal compartments. TBCK-deficiency leads to compartment-specific mRNA and lysosomal trafficking defects in neurons, which likely contribute to the preferential susceptibility to neurodegeneration.

## INTRODUCTION

The cellular mechanisms underlying selective neuronal vulnerability in neurodegenerative diseases remain poorly understood. In children, as opposed to adults, these disorders are often monogenic and ultra-rare. Similar to adult-onset disorders, though, there is a dire lack of disease-modifying therapies. We previously reported a pediatric onset disorder of variable severity associated with biallelic variants in the *TBCK* gene^1^. Due to a founder effect, many affected children are of Puerto Rican descent and share the *TBCK* “Boricua” (p.R126X) stop-gain homozygous variant. Clinically, we referred to the disorder as TBCK-encephaloneuronopathy (TBCKE) to highlight features not uniformly reported across all patients with biallelic variants in TBCK. TBCKE leads to progressive motor neuron degeneration, white matter lesions, and diffuse cortical atrophy. Clear genotype-phenotype correlations across *TBCK* variants remain to be established, as some patients have been reported with a milder phenotype^2^ and without clear evidence of progression.

*TBCK* gene encodes the protein <u>TBC</u>1-domain containing <u>K</u>inase^1^. Despite the disease’s primarily neurological presentation, *TBCK* mRNA and protein are expressed in most human tissues^3^. The physiologic function of TBCK protein is poorly understood. Knockdown of this protein is associated with down-regulation of mTORC1 signaling in HEK293 cells^4^. TBCKE patient-derived fibroblasts show evidence of aberrant autophagy^5^, and mitochondrial respiratory defects secondary to impaired mitochondrial quality control due to lysosomal dysfunction^6^. Neuropathological data also support a significant role for autophagic-lysosomal dysfunction in the pathophysiology of the disease, revealing the accumulation of intraneuronal lipofuscin storage material^7^.

There is growing evidence that mRNA can be trafficked along organelles, particularly in neurons. mRNA has been shown to traffic within axons together with late endosomes^8^ and lysosomes^9^. There is also recent evidence that mRNA can be transported in association with mitochondria, for instance, to support local mitophagy and fission^10,11^. The significance of these observations is that there is an emerging role for deficits in neuronal mRNA trafficking across various neurodevelopmental and neurodegenerative disease models. For instance, recent data in models of motor neuron disease (ALS) suggest that TDP-43 accumulation in axons ultimately disrupts ribonucleoprotein (RNP) assembly and trafficking, leading to inhibition of local protein synthesis in the distal axonal mitochondria of human iPSC-derived motor neurons^12^.

TBCKE patients also exhibit prominent motor neuron deficits and evidence of mitochondrial dysfunction. TBCK protein was recently found to be a component of a novel mRNA transport complex, denominated five-subunit endosomal Rab5 and RNA/ribosome intermediary (FERRY) complex^13^. Intriguingly, of the five proteins proposed to form the FERRY complex (TBCK, PP1R21, C12orf4, CRYZL1 and GATD1)^13^. three are linked to severe, pediatric-onset, neurologic disorders. Biallelic loss of function variants in PPP1R21 have been linked to a neurodevelopmental disorder with remarkable clinical overlap to TBCK^14,15^, just as biallelic variants in *C12orf4* cause neurodevelopmental delays and intellectual disability^16,17^. It remains unclear if there was clinical evidence of progression or neurodegeneration in the handful of patients reported. In the recently reported cryo-EM structure of the FERRY complex^18^, PPP1R21 is proposed as the predominant mRNA binding protein, which also serves as a hub connecting all complex subunits and mediates anchoring the complex to early endosomes via RAB5.

The structure of TBCK remains unresolved, yet the profound impact of loss of function variants in neurologic function implies that its physiological function is crucial for maintaining neuronal homeostasis. Most previous studies of TBCK-deficiency have used non-neuronal cell models, although one study examined TBCK function using iPSC-derived neural progenitor cells (iNPCs) from controls or patients with biallelic LOF variants of the *TBCK* gene (exon 23 deletion and p,Tyr710*)^19^. They found that TBCK colocalizes with proteins that regulate vesicular trafficking in the endocytic and secretory pathways (RAB5a, COPII and clathrin). Yet it did not find evidence of altered endosome formation or maturation in NPCs from *TBCK* patients. In non-neuronal cells, TBCK has been shown to modulate mTORC1 activation levels and colocalize with gamma-tubulin along the mitotic spindle^4^. To our knowledge, there are also no published animal models for loss of TBCK. Hence the molecular consequences of the loss of TBCK in neurons remain largely unexplored.

Herein we report that TBCK deficiency in patient-derived neurons results in significantly reduced neuronal survival and aberrant neuronal morphology. TBCK localizes to early endosomes but is also detected in proximity with LAMP2 (late endosomes/ lysosomes). Importantly, endolysosomal trafficking and/or maturation in neurons is disrupted by loss of TBCK expression, leading to perinuclear accumulation of enlarged early endosomes, and aberrant localization of the FERRY protein PPP1R21. We find a novel interaction with JIP4, an adaptor for the dynein/dynactin microtubule minus-end directed motor complex, across heterologous human cellular models, including HEK293, fibroblasts, lymphoblastoid cell lines (LCLs) and neurons. Our data also reveals that TBCK may regulate JIP4 levels in a tissue-specific fashion, as JIP4 levels are significantly reduced in TBCK-deficient neurons but not fibroblasts. Accordingly, we demonstrate that TBCK-deficiency leads to a dramatic defect of axonal lysosomal motility and reduced axonal mRNA content. Our data support a model in which TBCK-deficiency impairs trafficking/and or maturation of endosomes and lysosomes across cell types, and specifically in neurons, impairs axonal lysosomal retrograde transport.

## RESULTS

### TBCK in neurons localizes primarily to vesicular structures

After reprogramming patient-derived fibroblasts homozygous for the Boricua *TBCK* variant (c.376 C>T, p.R126X) to generate iPSCs, we utilized the neurogenin-2 (Ngn2) differentiation protocol^20^ to generate iPSC-derived neurons (iNeu) (Fig. 1a). iPSC derived from healthy donors were used as control lines. We first tested whether TBCK-deficient cells undergo differentiation by assaying expression of neuron-specific mRNAs and proteins. We found that the levels of *GRIN1, GRIN2A*, *GRIN2D*, and *GRIN3A* mRNA increased over time during differentiation similarly in control and TBCK-deficient neurons (assayed at days 3, 7 and 14) (Fig. S1a-d). Similarly, immunoblotting showed that neuronal marker beta-III tubulin (TUJ1) was expressed at similar levels in control and TBCK-deficient cells (Fig S1e-f). These results show that TBCK-deficiency due to homozygous p.R126X variant does not grossly impact the neuronal differentiation potential of iPSC via neurogenin induction.

**Fig. 1:**
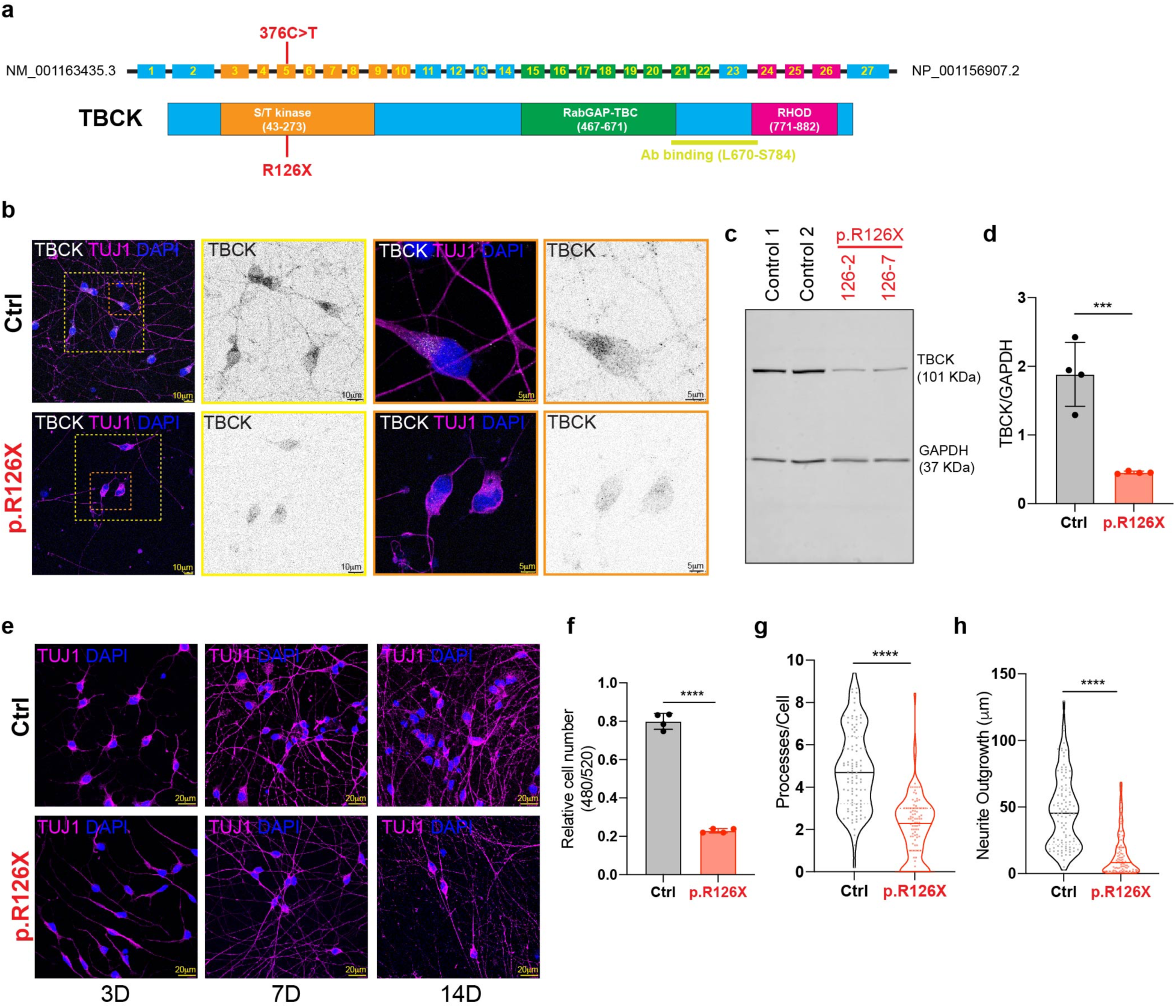
p.R126X mutation causes TBCK deficiency, morphological alterations and reduces neuronal survival. **a,** Schematic representation of TBCK gene and protein. Mutation p.R126X (376C>T) is indicated relative to their position in the DNA. **b,**TBCK detection by immunofluorescence in iPSC-derived neurons (iNeurons) at 14 days of differentiation (14D) in control (Ctrl) and TBCK-deficient cells (p.R126X). **c,** Immunoblotting analysis of TBCK levels in control and p.R126X iNeurons (14D). **d,** Quantification of TBCK protein levels relative to GAPDH (n=4). Significance was calculated using unpaired t-test analysis. **e,** Immunofluorescence of TUJ1 showing morphologies of iNeurons at 3, 7 and 14D. **f,** Quantification of cell number by detention of nucleus signal employing CyQUANT (n=4). Significance was calculated using unpaired t-test analysis. **g,** Quantification of neuronal processes in control and p.R126X iN (n=3). Significance was calculated using unpaired t-test analysis. **h,** Determination of neurite outgrowth in control and p.R126X iN (n=3). Significance was calculated using unpaired t-test analysis. ***p<0.001 and ****p<0.001.All graphs show error bars with mean ± SD from independent experiments (n).

We then assayed TBCK mRNA levels by RT-PCR during differentiation in control and patient-derived cells at day 3, 7, and 14. Interestingly, we saw TBCK expression levels increase in control cells as they progressed to neuronal differentiation, with significantly higher levels at day 14 compared to day 3 (Fig. S1g). This correlated with protein levels measured by immunoblot (Fig. S1h). Unexpectedly, we also found detectable *TBCK* mRNA levels in patient-derived iNeu, suggesting a degree of either escape of non-sense mediated decay or alternative splicing leading to skipping the nonsense variant ^21^. Indeed, examining RNA transcript databases, we found evidence that an alternative transcript where exon 5 (including c.376 C>T) is spliced out, is the second highest most expressed transcript (after the canonical transcript and present in human brain (https://gtexportal.org). Hence throughout this manuscript, we refer to patient-derived neurons as TBCK-deficient, as some trace levels of protein and mRNA were detectable.

We then queried the subcellular localization of TBCK protein in our human iPSC-derived models. Previous literature suggests TBCK protein can localize to multiple subcellular compartments across various cell types, ranging from diffuse cytoplasmic, nuclear, associated with the mitotic apparatus, and vesicular^4,19,21^. Yet the subcellular localization of TBCK in neurons has not been previously reported. Therefore, we assayed TBCK protein subcellular localization by immunostaining in control and patient-derived iNeu at day 14. TBCK was detected in control iNeu in both the soma (including low levels in nucleus) and neuronal processes, in a predominantly punctate pattern, suggesting localization to a cytoplasmic organelle. Consistent with low level of mRNA transcripts detected in patient-derived iNeu, trace signal was detectable by immunostaining in p.R126X iNeu (Fig. 1b). Immunoblotting confirmed that patient-derived iNeu have significantly reduced levels of full-length TBCK protein relative to control iNeu (24% of control) (Fig. 1c, d). Overall our data show that in human neurons TBCK protein subcellular localization is primarily vesicular, although nuclear staining is also detectable in variable levels.

### TBCK-deficiency in patient-derived neurons leads to reduced survival and aberrant morphology

Despite the similar expression of early neuronal differentiation markers in control and patient-derived iNeu, we found that TBCK-deficient iNeu exhibit reduced and limited viability beyond day 14 of differentiation. At this time point, TBCK-deficient iNeu grossly had smaller cell bodies and thinner projections (Fig 1e and S1i). Despite equal initial plating density, TBCK-deficient iNeu had significantly reduced cell counts relative to controls at day 14 control iNeu (Fig. 1f). Quantification by high content confocal imaging at day 14 showed that TBCK-deficient iNeu had fewer processes per cell (Fig. 1g) and neurites (Fig. 1h) than control iNeu. Therefore, we find that TBCK-deficiency in human iPSC-neuronal models leads to aberrant morphology and reduced survival.

### TBCK-deficient neurons recapitulate fibroblast mitochondrial findings but suggest cell-type specific effects on mTORC1 signaling

TBCK knockdown was previously reported to lead to reduced cell size and proliferation due to mTORC1 signaling deficits in HEK293 cells^4^. In patient-derived fibroblasts, we also found evidence of increased autophagic flux and mitochondrial respiratory defects^5^. To determine if these findings extend to neurons, we compared markers for the mTORC1 pathway, constitutive autophagy, and mitochondrial respiratory capacity in control versus TBCK-deficient iNeu.

TBCK-deficient iNeu (day 14) had no significant changes in total mTOR protein levels by immunoblot and a small (15%) but statistically significant decrease of phospho-mTOR levels (Fig. 2a,b), suggesting a mild reduction in activity of mTORC1 signaling. In terms of autophagy, TBCK-deficient iNeu expressed higher levels of LC3-II (Fig. 2a,c) and lower levels of SQSTM1/p62 (Fig. 2d and 2e), suggesting increased levels of baseline autophagy. These data suggest that neurons recapitulate the upregulation of autophagy seen in fibroblast models, but that effects on mTORC1 signaling in neurons were not as substantial as in other cellular models (36% reduction in lymphoblasts and >50% reduction in HEK293 cells)^2,4^.

**Fig. 2:**
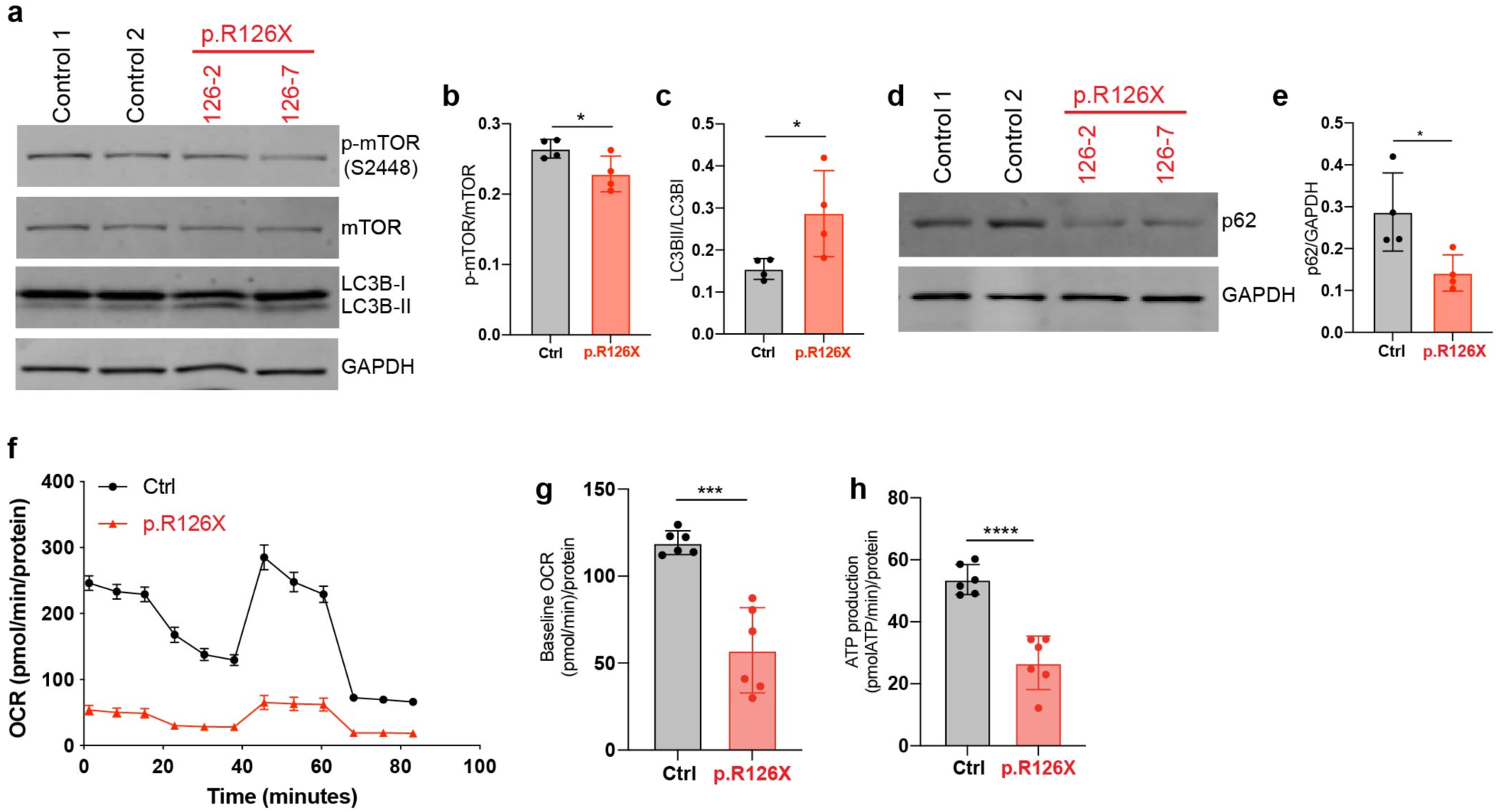
TBCK-deficient neurons show increased autophagy flux and alterations in mitochondrial respiration. **a,** Immunoblot of mTOR, phospho-mTOR (p-mTOR) and its downstream target LC3B in iNeurons at 14D in control (Ctrl) and TBCK-deficient neurons (p.R126X). **b-c,** Quantification of p-mTOR and LCB-II in relative to their total levels (n=4). Significance was calculated using unpaired t-test analysis. **d,** Immunoblot of p62 in control (Ctrl) and TBCK-deficient neurons (p.R126X) at 14D. **e,** Quantification of p62 protein levels relative to GAPDH (n=4). Significance was calculated using unpaired t-test analysis. **f,** Seahorse assay showing mitochondrial Oxygen Consumption Rate (OCR) (n=6). **g,** OCR values corresponding to mean for each cell line baseline respiration (n=6). Significance was calculated using unpaired t-test analysis. **h,** Metabolic flux analysis showing mitochondrial ATP production rate (n=6). Significance was calculated using unpaired t-test analysis. *p<0.01, ***p<0.001 and ***p<0.001. All graphs show error bars with mean ± SD from independent experiments (n).

We then assayed mitochondrial function by Seahorse assays and found that relative to control iNeu, TBCK-deficient iNeu have significant impairment in oxidative phosphorylation, as evidenced by reduced oxygen consumption rate (OCR) (Fig. 2f,g) and ATP production (Fig. 2h). Immunoblotting did not detect significant differences between control and TBCK-deficient iNeu in total levels of mitochondrial content such as outer membrane protein TOM20 (Fig. S2a,b) or subunits of mitochondrial respiratory chain complexes (Fig. S2c-i). These data show that respiratory defects seen in TBCK-deficiency are not due to a significant reduction in mitochondrial content or electron transport chain complex subunits.

### Unbiased neuronal proteomics reveal pathways altered in TBCK-deficient neurons linked to common neurodegenerative disorders

To further explore the molecular consequences of TBCK deficiency in neurons, we performed a comparative proteomic analysis in controls versus TBCK-deficient iNeu at 14 days. In total, 6001 proteins were identified by mass spectroscopy, with 13 proteins detected only in control neurons and 10 only in TBCK-deficient neurons (Fig. S3a). Of the 6001 detected proteins, 894 were significantly downregulated (p<0.05) and 680 upregulated (p<0.05) in TBCK-deficient neurons relative to controls (Fig. 3a). Using log2 fold change >1 (corresponding to a 2-fold change in protein expression) and p<0.05 as filtering criteria, Gene ontology (GO) enrichment analysis ^22^ was performed. Significantly upregulated proteins in TBCK-deficient iNeu were primarily associated with neuronal projection development and synapse organization, as well as vesicles and their transport (Fig. 3b and S3b). KEGG pathway analysis identified “Neurodegeneration” as the most significant pathway, and in the top 10 significant pathways identified, Parkinson’s, Huntington’s and Alzheimer’s disease were also found (Fig. 3c and S3c). On the other hand, in terms of downregulated proteins, GO analysis shows an enrichment of proteins associated with nuclear processes (Fig. S3d-f).

**Fig. 3:**
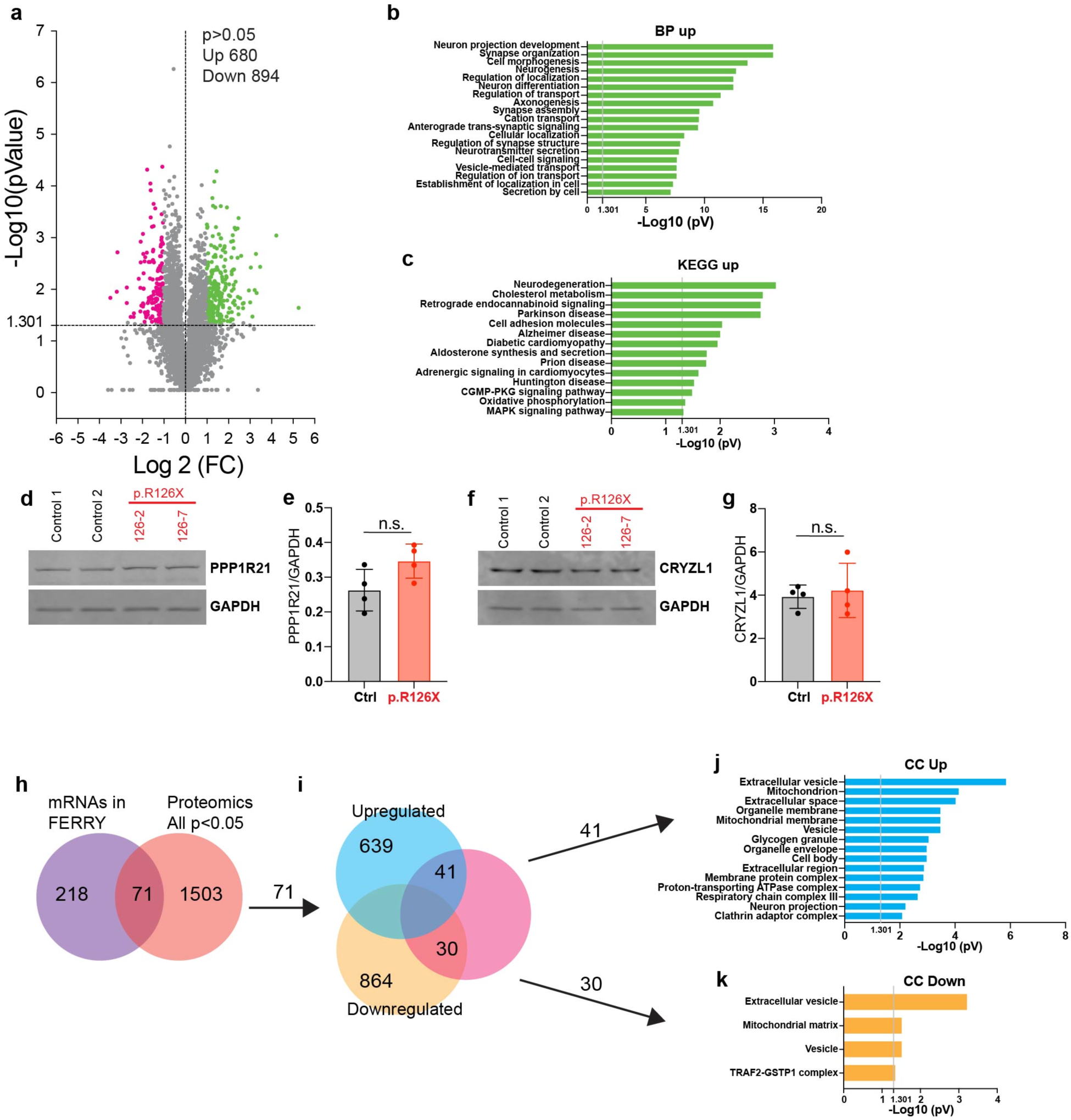
Proteomics in TBCK-deficient neurons reveals alterations in vesicular transport and associations with neurodegenerative diseases. **a**, Volcano plot showing changes in abundance of proteins detected in iNeurons p.R126X compared with controls. Dashes lines indicate statistical significance cut off (horizontal, p<0.05 or -log10>1.301) and direction of fold change (vertical, left and right quadrants denote decreased and increased abundance, respectively (n=4). **b,** Gene Ontology (GO) classification of up-regulated proteins in TBCK-deficient neurons based on biological process (BP) (cut off p<0.05 and FC>1). **c,** Pathway analysis of up-regulated proteins, according to the Kyoto Encyclopedia of Genes and Genomes (KEGG) database (cut off p<0.05 and FC>1). **d-g,** Levels of proteins (PPP1R21 and CRYZL1) in iNeurons (controls and p.R126X) (n=4). **h,** Comparison between mRNAs bound to FERRY complex and our proteomics data. Significance was calculated using unpaired t-test analysis. **i,** Venn diagram showing upregulated proteins (41) or downregulated (30) from filtered mRNAs (71). **j,** GO classification based on cellular component (CC) of upregulated proteins from mRNAs (41) found in the FERRY complex. **k,** Classification by cellular component (CC) of downregulated proteins from mRNAs (30) found in the FERRY complex. Significantly enriched GO terms are shown with Benjamini-Hochberg FDR-corrected p-values. All graphs show error bars with mean ± SD from independent experiments (n).

We then asked if TBCK-deficient neurons exhibit significant changes in the levels of the proteins that form the FERRY (Figure S3h-i) complex (PPP1R21, CRYZL1 and GATD1) by proteomics. Reassuringly, TBCK was amongst the proteins that were only detected in control but not in patient-derived iNeu by mass spectroscopy. We found no significant changes in the levels of CRYZL1 and GATD1 (Fig. S3g,h) and only a slight increase in PPP1R21 levels (Fig. S3i). These results were validated using immunoblots and found to be consistent across all the FERRY complex proteins in control vs TBCK-deficient iNeu (Fig. 3d-g).

Finally, we sought to test if we could detect in our proteomics data set differences in levels of proteins that are encoded by the mRNAs previously reported to interact directly with the FERRY complex^13^. Of the 1574 proteins differentially expressed in our dataset, we identified 71 proteins encoded by mRNAs associated with FERRY (289) (Fig. 3h,i). Both the upregulated (41) and downregulated (30) proteins in our dataset were largely related to vesicles and mitochondria (Fig. 3j,k). Therefore, proteomic analysis did not detect distinct up or downregulation of specific pathways when examining the subset of proteins encoded by transcripts linked to the FERRY complex.

### Neuronal protein interactome identifies novel interactors of TBCK and validates TBCK as a member of the FERRY complex

To gain further insight into the physiologic function of TBCK in a disease-relevant model, we performed immunoprecipitation of TBCK followed by mass spectrometry to determine a protein-protein interactome of TBCK. We identified 37 potential TBCK interactors, and obtained their putative localization and functions using existing databases (Human Protein Atlas (https://www.proteinatlas.org) and GeneCards (https://www.genecards.org) (Fig. S4a). Thirteen (13) of the 37 proteins interacting with TBCK (PPP1R21, CRYZL1, C12orf4, TRI27 (TRIM27), JIP4, MED23, SYYM, FSBP, CEP97, ODFP2, DAGLB, MMAB and SFXN1) were considered high confidence interactors (Fig. 4a) based on the following criteria: i) minimum absolute fold change of 2 (Immunoprecipitated with anti-TBCK versus anti-IgG), ii) q-value <0.1, iii) identified by a minimum of 2 razor+ unique peptides in any sample, and iv) detected in at least 2 of the experimental replicates. Amongst these high-confidence interacting proteins, PPP1R21, TRIM27 and JIP4, have been previously associated with endolysosomal vesicular pathways^15,23,24^. PPP1R21 has been associated with both early endosomes and more recently the FERRY complex. While JIP4 functions as a microtubule motor adaptor, TRIM27 (Tripartite motif-containing protein 27, also called RFP) is proposed to function as a multifunctional ubiquitin E3 ligase implicated in multiple cellular processes including endosomal trafficking and mitophagy^25^. Besides proteins linked to vesicular trafficking, additional novel interactors identified suggest that TBCK protein may have relevant interactions across other subcellular compartments, including the centrosome/microtubule organizing center (ODFP2, CEP97), the nucleus (MED23 and FSBP) and mitochondria (MMAB, SFXN1 and SYYM).

**Fig. 4:**
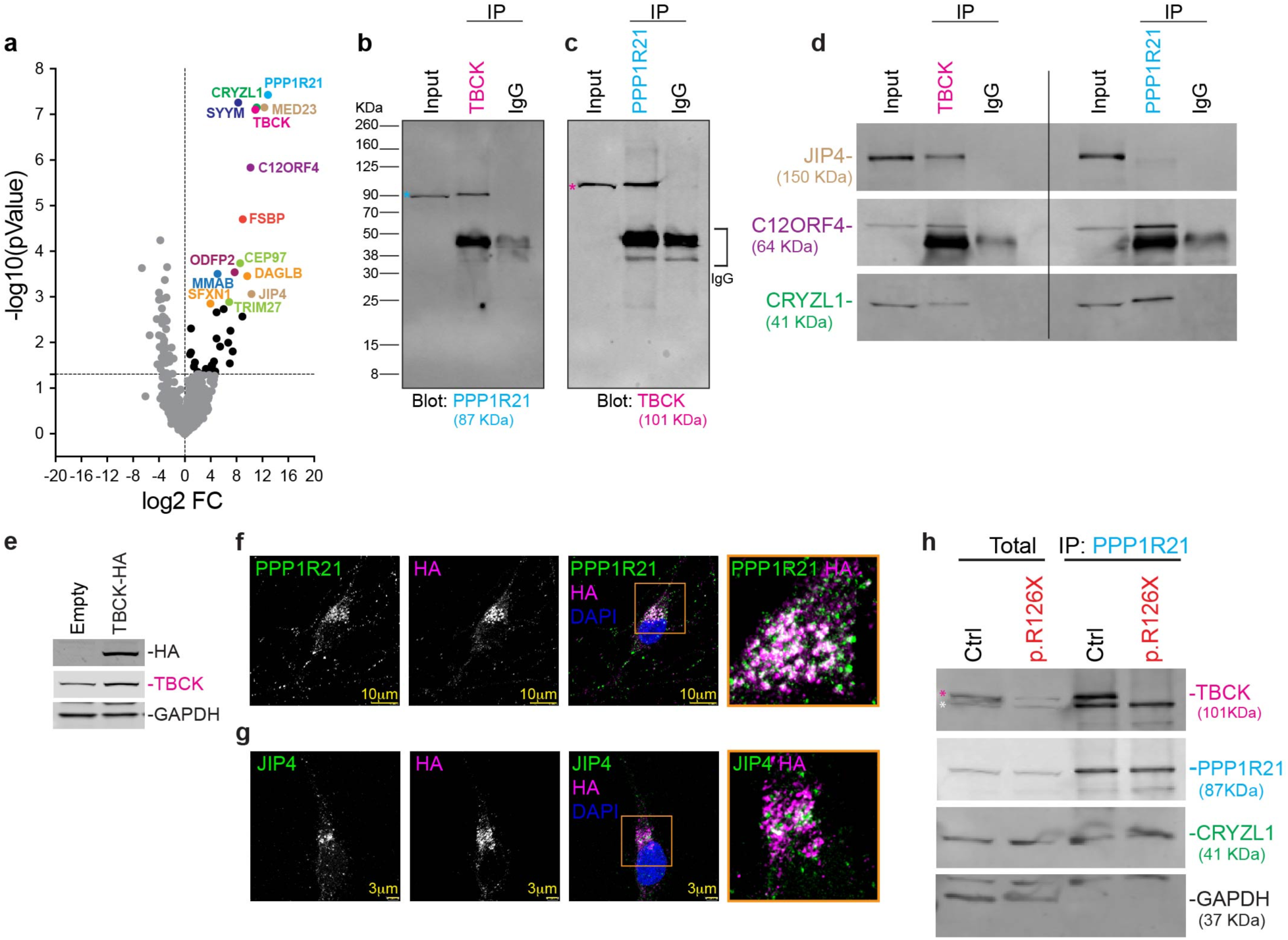
TBCK interactome shows novel interactions associated with the FERRY complex. **a,** Volcano plot showing mass spectrometry of proteins pulled down with TBCK performed in control iNeurons. The plot shows the log 2-difference in abundance of each protein with cut off p<0.05 (or -log10> 1.301). Interactors with high confidence are showed in colors different to black or gray (n=3). **b,** Representative immunoblot of PPP1R21 (blue asterisk) co-immunoprecipitated with TBCK in iNeurons. Input was loaded with 20μg of proteins. **c,** Immunoblot of TBCK (pink asterisk) co-immunoprecipitated with PPP1R21. Input was loaded with 20μg of proteins. **d** Left panel: Immunoblot of proteins co-immunoprecipitated with TBCK in iNeurons. Right panel: Immunoblot of proteins co-immunoprecipitated with PPP1R21. Input was loaded with 20μg of proteins. **e,** Immunoblot of iPSC-derived neurons expressing TBCK-HA or not (empty). **f,** Representative immunostaining of HA and PPP1R21 in iPSC-derived neurons expressing TBCK-HA. Orange square shows image magnification. **g,** Immunostaining of HA and JIP4 in iPSC-derived neurons expressing TBCK-HA. Orange square shows image magnification. **h,** Immunoblot of immunoprecipitated PPP1R21 from control and TBCK-deficient neurons (p.R126X). Pink asterisk identifies TBCK band and white PPP1R21.

Our interactome data is consistent with an interaction of TBCK with members of the recently proposed FERRY complex, namely PP1R21, CRYZL1 and C12ORF4, which had been previously identified as a single complex through affinity purification of RAB5 effectors from bovine brain ^13^. However, our interactome did not detect TBCK interactions with GATD1, which was also previously reported as a FERRY complex component ^18^. We further validated the interaction of TBCK and PPP1R21 by co-immunoprecipitation (Co-IP), showing that TBCK and PPP1R21 co-immunoprecipitate regardless of whether TBCK or PPP1R21 was used as bait (Fig. 4b,c). We also validated the interactions of C12ORF4, CRYZL1 TBCK, and PPP1R21 both in neurons (Fig. 4d) and HEK293 cells expressing Flag-TBCK by Co-IP (Fig. S4b). In order to further test the specificity of these novel interactions, we also generated TBCK-HA iNeu (Fig.4e). Immunostaining for HA and FLAG demonstrated robust colocalization with PPP1R21 (Fig, 4f and Fig S4c) in both cell types, and TRIM27 (Fig. S4d) in HEK cells. Overall, our data supports that TBCK forms stable interactions consistent with the proposed FERRY protein complex, with the notable exception of GATD1, in various human cellular models.

### TBCK interacts with JIP4, linking the FERRY complex to the microtubule motor machinery

Interestingly, our neuronal interactome data identified a novel interaction of TBCK with the microtubule motor adapter protein, JIP4. We then asked whether JIP4 interacts with all members of the FERRY complex or exclusively with TBCK in various human cellular models. While immunoblotting for JIP4 after the pulldown of TBCK validated the interaction with JIP4, we found no evidence of interaction with JIP4 when using PPP1R21 as bait (Fig. 4d). We also tested the interactions of TBCK with JIP4 and other members from the FERRY complex (PP1R21, C12ORF4) using human fibroblast and Lymphoblastoid Cell Lines (LCLs) (Fig. S4e). In all cell lines, we found that TBCK coimmunoprecipitates with JIP4. Lastly, to further confirm this interaction and control for any potential non-specific antibody binding, we tested the TBCK-Flag in HEK293 cells with the anti-TBCK antibody (Fig S4f). Neurons expressing TBCK-HA show colocalization of TBCK with JIP4 (Fig. 4g). In HEK cells immunostaining for anti-Flag and JIP4 demonstrated strong colocalization of TBCK with JIP4 (Fig. S4g). Taken together, our data is consistent with JIP4 specifically interacting with TBCK and not with other complex members like PP1R21. Hence, this suggests the TBCK-JIP4 interaction may play a role in linking the FERRY complex to microtubule motor machinery to mediate intracellular trafficking.

### TBCK-deficiency differentially modulates levels of interactors JIP4, TRIM27 and C12orf4 across cell types

To further understand the downstream consequences of TBCK-deficiency in patient derived neurons, we then examined how loss of TBCK affected the levels of its known interactors, including the members of the FERRY complex. We found that TBCK-deficient neurons show no significant difference versus controls in levels of PPP1R21 and CRYZL1 (Fig. 4h). We also found that CRYZL1 was detected at similar levels in PPP1R21 immunoprecipitates from control and TBCK-deficient neurons (Fig. 4h). These experiments suggest that the presence of TBCK is not necessary for the interaction of PPP1R21 with CRYZL1 in human neurons.

We also examined the TBCK interactors in our proteomics data set, including the members of the FERRY complex (Fig. S5a). This revealed that the levels of TBCK-interacting proteins C12ORF4, JIP4 and TRIM27 were significantly reduced in TBCK-deficient iNeu relative to controls (Fig. S5b-d). Immunoblotting experiments confirmed that TBCK-deficient neurons show a drastic reduction in levels of JIP4, C12orf4, and TRIM27 (Fig. 5a-d). Interestingly, we found no difference between control and TBCK-deficient neurons in mRNA expression levels of *C12orf4, SPAG9* (encoding JIP4) or *TRIM27* (Fig. 5e-g). Hence our data suggest, that post-translational (the translation itself may also be affected) mechanisms mediate the loss of these proteins in TBCK-deficient neurons.

**Fig. 5:**
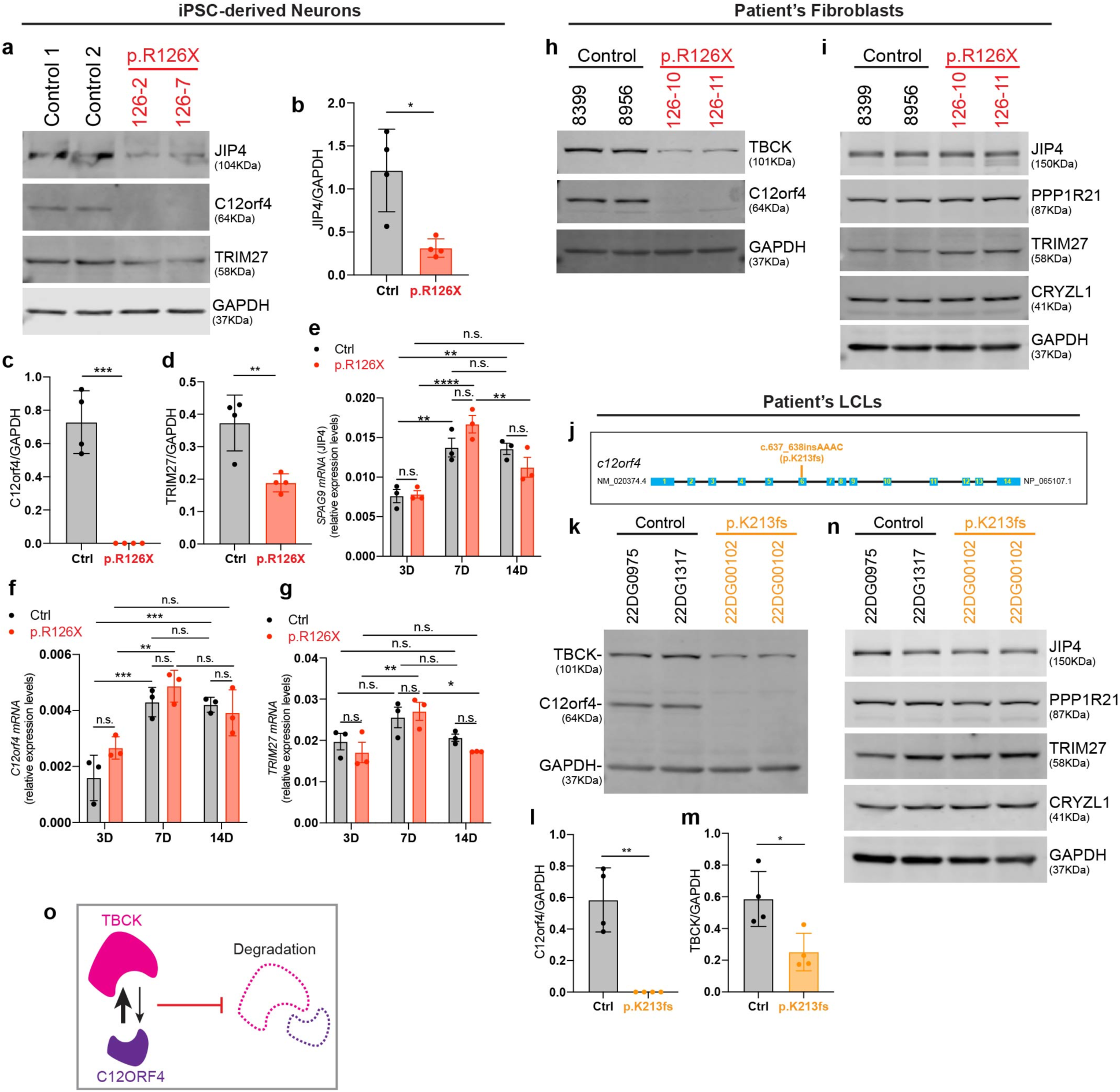
TBCK post-transcriptionally regulates JIP4, C12orf4 and TRIM27 in cell-type-specific fashion. **a,** Representative immunoblot showing less proteins levels of JIP4, C12orf4 and TRIM27 in controls (Ctrl) and TBCK-deficient neurons (p.R126X). **b,** Quantification of protein levels of JIP4, (**c**) C12orf4 and (**d**) TRIM27 in controls (Ctrl) and TBCK-deficient neurons (p.R126X) (n=4). Significance was calculated using unpaired t-test analysis. **e,** mRNA expression of *SPAG9* (JIP4), (**f**) C12orf4 and (**g**) *TRIM27* at different stages of differentiation (3, 7 and 14 days) in controls (Ctrl) and TBCK-deficient neurons (p.R126X) (n=3). Significance was calculated using two-way ANOVA with Sidak’s post hoc analysis for multiple comparison. **h,** Representative immunoblot of TBCK and C12orf4 in control and TBCK-deficient fibroblasts (p.R126X). **i,** Immunoblot of JIP4, PPP1R21, TRIM27, and CRYZL1 in control and TBCK-deficient fibroblasts (p.R126X). **j,** Schematic representation of C12orf4 gene showing the variant p.K213fs (c.637_638insAAAC) relative to its position in the DNA. **k,** Representative immunoblot showing deficiency of TBCK and C12orf4 in affected lymphoblastoid cell lines (LCLs) (p.K213fs) compared to controls. **l,** Quantification of protein levels of C12orf4 and (**m**) TBCK in control (Ctrl) and affected LCLs (p.K213fs) (n=4). Significance was calculated using unpaired t-test analysis. **n,** Immunoblots of C12orf4 and its interactors (JIP4, PPP1R21, TRIM27, and CRYZL1) in TBCK in control (Ctrl) and affected LCLs (p.K213fs). **o,** TBCK and C12ORF4 mutually regulate their protein levels, being TBCK the main regulator in this interaction. *p<0.01, **p<0.05, ***p<0.001 and ****p<0.001. All graphs show error bars with mean ± SD from independent experiments (n).

We then asked if TBCK deficiency modulated levels of its interactors similarly across different human cell types. We found that TBCK deficiency consistently leads to a robust reduction of C12orf4 across all cell models tested, whether primary patient-derived fibroblasts (Fig. 5h,i) or in control cell lines (HEK293 and iPS cells) where TBCK was acutely knocked down using shRNA (Fig. S5e). On the contrary, levels of JIP4 and TRIM27 were significantly reduced in neurons (Fig. 5a) but not in fibroblasts (Fig. 5i). Consistent with the fibroblast data, shRNA-mediated TBCK knockdown did not affect JIP4 or TRIM27 levels (Fig. 5e). Therefore, these results reveal TBCK may regulate levels of interactors in a tissue-specific fashion, with depletion of JIP4 and TRIM27 occurring as a neuronal specific phenotype.

### C12orf4 and TBCK mutual interaction is relevant to human disease

Biallelic variants in the gene *C12orf4* cause a severe neurodevelopmental disorder (MIM# 618221) with striking clinical similarities to TBCK encephaloneuronopathy^16^. We assayed cells (LCLs) from a patient homozygous for loss of function variants (p.K213fs) in *C12orf4*^16,26,27^ (Fig. 5j). We confirmed that indeed C12orf4 protein levels were depleted (Fig. 5k,l) in patient LCLs (as previously reported by Alazami^16^, LCLs were a gracious gift from Dr. Alkuraya). Given our data showing that loss of TBCK may destabilize C12orf4 protein, we asked if loss of C12orf4 expression would also affect TBCK protein levels. Interestingly, we found that TBCK protein levels were reduced by 43% in patient cells compared to parental control LCLs (Fig. 5k,m).

On the other hand, immunoblot for FERRY complex members PPP1R21 and CRYZL1 showed these proteins were unaffected by loss of C12orf4 (Fig. 5n and S5f,g). We also found no significant difference in the levels of TBCK interactors JIP4 or TRIM27 in the C12orf4 patient’s LCLs (Fig. 5n and S5h-i). Our results show that while C12orf4 stability is highly dependent on TBCK, loss of C12orf4, at least on LCLs, leads to a significant decrease in TBCK protein levels but not the other FERRY complex proteins. Hence, the mutual regulation between TBCK and C12orf4 occurs at a post-transcriptional level, TBCK may play a role in stabilizing/regulating C12orf4 (Fig. 5o).

### TBCK deficiency alters early endosome subcellular localization and distribution

Our proteomics and interactome data strongly suggest that TBCK physiologic function impacts cellular processes related to the endolysosomal pathway. Hence, we then examined how TBCK deficiency may affect early endosomes in our cellular models. We performed immunostaining for early endosomes (using RAB5 and EEA1 as markers) in control and TBCK-deficient iNeu. We found that both RAB5+ (Fig. 6a and S6a) and EEA1+ vesicles (Fig. 6b and S6b) were preferentially located in the soma and/or perinuclear area of TBCK-deficient iNeu compared to controls. We consistently found this phenotype of perinuclear enlarged vesicles in patient-derived neurons using three different early endosome markers (EEA1, RAB5, and PPP1R21) (Fig. 6c,d).

**Fig. 6:**
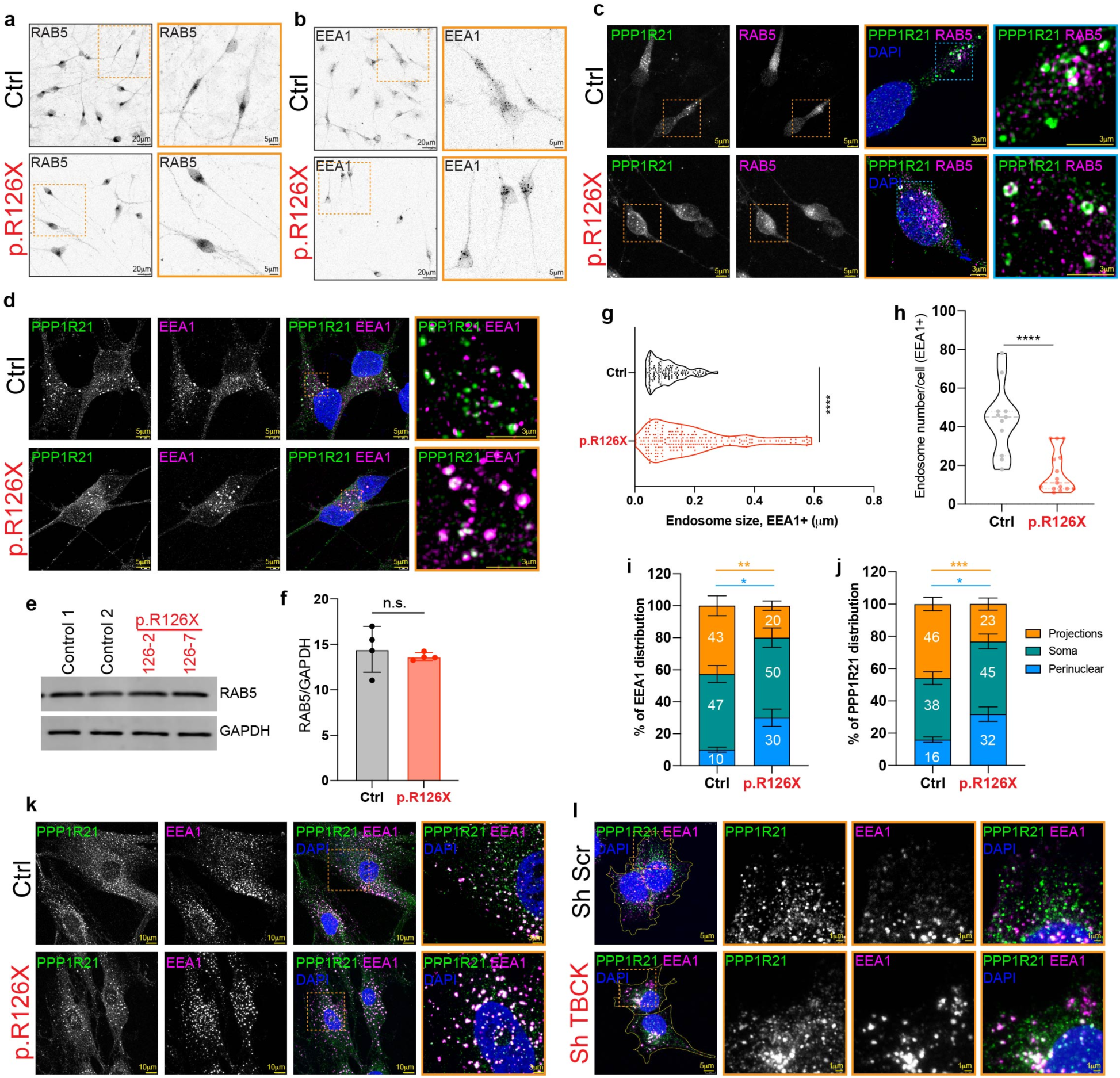
TBCK regulates the distribution and size of early endosomes through PPP1R21. **a,** Representative confocal images showing RAB5 and (**b**) EEA1 in control (Ctrl) TBCK-deficient neurons (p.R126X). Inset magnifications (orange squares) show the distribution of early endosomes. **c,** Confocal images show PPP1R21 and RAB5 in control (Ctrl) TBCK-deficient neurons (p.R126X). Different inset magnifications (orange and blue squares) show the distribution and colocalization of PPP1R21 and RAB5. **d,** Confocal images showing PPP1R21 and EEA1 in control (Ctrl) TBCK-deficient neurons (p.R126X). Different inset magnifications (orange and blue squares) show the distribution and colocalization of PPP1R21 and EEA1. **e,** Representative immunoblot and (**f**) quantification of the early endosome marker, RAB5, in control (Ctrl) TBCK-deficient neurons (p.R126X). Significance was calculated using unpaired t-test analysis. **g,** Analysis of endosome size showing larger endosomes in TBCK-deficient neurons (p.R126X) compared to controls (Ctrl) (n=3). Significance was calculated using unpaired t-test analysis. **h,** Quantification of endosomes number shows fewer number of endosomes (EEA1+) in TBCK-deficient neurons (p.R126X) compared to controls (Ctrl) (n=3). Significance was calculated using unpaired t-test analysis. **i,** EEA1 and (**j**) PPP1R21 distribution along compartments of control (Ctrl) TBCK-deficient neurons (p.R126X). Cell compartments were identified and traced using Fiji, ImageJ (n=3). Significance was calculated using unpaired t-test analysis. **k,** Confocal images show PPP1R21 and EEA1 in control (Ctrl) TBCK-deficient fibroblasts (p.R126X). Different inset magnifications (orange squares) show the distribution and colocalization of PPP1R21 and EEA1. **l,** Staining of PPP1R21 and EEA1 in HEK293 cells transduced with ShRNA scramble (Sh Scr) and ShRNA TBCK#2 (Sh TBCK). Different inset magnifications (orange squares) show the distribution and colocalization of PPP1R21 and EEA1. *p<0.01, **p<0.05 and ***p<0.001. All graphs show error bars with mean ± SD from independent experiments (n).

As Rab5 and its effectors are known to be crucial for endosomal trafficking and maturation, we tested if endosome localization differences could be mediated by changes in Rab5 expression levels. We quantified total RAB5 protein levels by immunoblots in control and TBCK-deficient neurons and found no significant difference (Fig. 6e,f), suggesting TBCK deficiency may alter RAB5 activity and/or modulate downstream effectors. Quantification of puncta size confirmed that EEA1+ early endosomes were significantly larger in TBCK-deficient cells compared to controls (Fig. 6g). Despite the larger size of endosomes, we found a significantly reduced number of EEA1+ vesicles in TBCK-deficient neurons (Fig 6h).

To further characterize how TBCK-deficiency may impact early endosomes distribution specifically in neurons, we quantified early endosomes in distinct neuronal compartments. EEA1+ and PPP1R21+ vesicles were quantified in perinuclear, somatodendritic and axonal regions. TBCK-deficient neurons have significantly more early endosomes (both by EEA1+ and PPPR21+) in the soma and a relative paucity of early endosomes in neuronal processes relative to controls (Fig. 6i,j). Altogether, our data reveals that TBCK deficiency leads to aberrant subcellular distribution across endosomal vesicles expressing multiple markers (EEA1+, Rab5+, PPP1R21+). Specifically, in human neuronal we find significantly reduced early endosomes specifically in neuronal processes relative to the soma.

### TBCK-deficiency disrupts endolysosomal maturation and alters the subcellular distribution of FERRY protein PPP1R21

We examined the localization of PPP1R21 as it has been previously reported to localize to early endosomes^15^, but also due to its proposed function as the scaffold of the FERRY complex, linking the complex to early endosomes via its interaction with RAB5^13,18^. We found that in control iNeu, PPP1R21 is clearly present in RAB5+ vesicles but not exclusively (Fig. 6c), with some diffuse cytoplasmic staining also evident, consistent with previous data^15^. By contrast, we found PPP1R21 in TBCK-deficient neurons localizes predominantly to enlarged perinuclear RAB5+ (Fig. 6c) or EEA1+ (Fig. 6d) vesicles. This suggests TBCK may modulate the localization/trafficking of PPP1R21, and perhaps the FERRY complex, in early endosomes.

In order to test if TBCK can consistently alter the distribution of PPP1R21 across cell types, we also assayed patient’s fibroblast and HEK293 (TBCK knockdown) cells. Similar to neurons, TBCKE patient-derived fibroblasts also showed increased colocalization of PPP1R21 within EEA1+ vesicles. Furthermore, these vesicles are preferentially located in the perinuclear area compared to control fibroblasts, suggesting aberrant maturation or trafficking (Fig. 6k and S6c). TBCK shRNA knockdown HEK293 cells also showed that PPP1R21 preferentially colocalizes with large and perinuclear early endosomes in the setting of acute TBCK-deficiency (Fig. 6l).

We further asked if PPP1R21 could colocalize with vesicles that have matured beyond early endosomes, including late endosomes/lysosomes (LAMP2+ vesicles). We found that PPP1R21 indeed can colocalize with a subset of LAMP2+ vesicles across multiple (fibroblasts, iNeurons, HEK293) cellular models (Fig. S7a-c). While a subset of LAMP2+ vesicles (∼25 %) were also positive for PPPP1R21 in control fibroblasts, TBCK-deficient fibroblasts had a robust increase (∼87%) in vesicles co-expressing these markers (Fig. S7d). As expected, based on its previously reported localization in early endosomes, we found that PPP1R21 has a considerable colocalization with EEA1 in control fibroblasts (∼55%). Nevertheless, in the setting of TBCK deficiency, most EEA1+ vesicles are also PPP1R21+ (∼90%), suggesting loss of TBCK may lead to PPPR21 remain anchored aberrant endolysosomal vesicles (Fig. S7d). We also observed a robust phenotype of large perinuclear LAMP2+ vesicles across all three TBCK-deficient cell models (Fig S7a,b).

Overall, our data reveal a novel subcellular localization for PPP1R21 in lysosomes in control cells, expanding the concept that PPP1R21 may have functions beyond the early endosome^15^. Also, we show that across cellular models, TBCK-deficient cells consistently showed a predominance of enlarged, perinuclear endolysosomal vesicles expressing multiple typically minimally overlapping markers ^28^. Therefore, TBCK may also play a pivotal role in endolysosomal maturation and/or trafficking.

### FERRY complex proteins localize beyond early endosomes to late endosomes/lysosomes

Our results showing that not all PPP1R21+ vesicles were early endosomes (EEA1+) in control cells, and that PPP1R21 can be found colocalized to a subset of LAMP2+ vesicles, suggested that the FERRY complex could perhaps localize beyond early endosomes as they mature into lysosomes. In addition, previous works by us and others have shown that TBCK-deficiency leads to lysosomal dysfunction^6,7^. Therefore, we hypothesized that TBCK, as an additional marker for the FERRY complex, can also localize to LAMP1+ or LAMP2+ vesicles (late endosomes/ lysosomes) in human neurons.

We first examined localization of TBCK and LAMP1 (Figure 7a). We found TBCK in close proximity, and often on the surface, of LAMP1-positive vesicles in neuronal cell bodies. To test if lysosomal dysfunction previously reported was due to grossly reduced lysosomal content, we assessed total levels of LAMP1 (glycosylated and non-glycosylated) by immunoblot and found similar levels in control vs mutant neurons (Figure 7c-d). Both antibodies we used for FERRY markers (PPP1R21 and TBCK) were raised in the same host, making co-IP and colocalization studies not feasible. Hence, to further test the subcellular localization of TBCK in the context of the FERRY complex, we generated an TBCK-HA lentiviral construct. We then transduced control iPSCs and assayed the localization of HA-TBCK in relation to FERRY protein PPP1R21 and LAMP1. We found that HA-TBCK and PPP1R21 were predominantly colocalized in the soma (Figure 7b), validating that functionally the recombinant HA-TBCK is likely being integrated into the FERRY complex. We also found evidence of LAMP1+ and HA+ vesicles, confirming our hypothesis that TBCK, and likely FERRY, can indeed localize to lysosomes in control neurons. Surprisingly, we found that TBCK-HA and LAMP1 colocalization was more frequently and clearly colocalized in axonal compartments relative to the soma (Figure 7b). Overall, these data, taken together with the novel interaction with lysosomal adapter protein JIP4, reveal that TBCK plays a role in localization of the FERRY complex to axonal lysosomes.

**Fig. 7:**
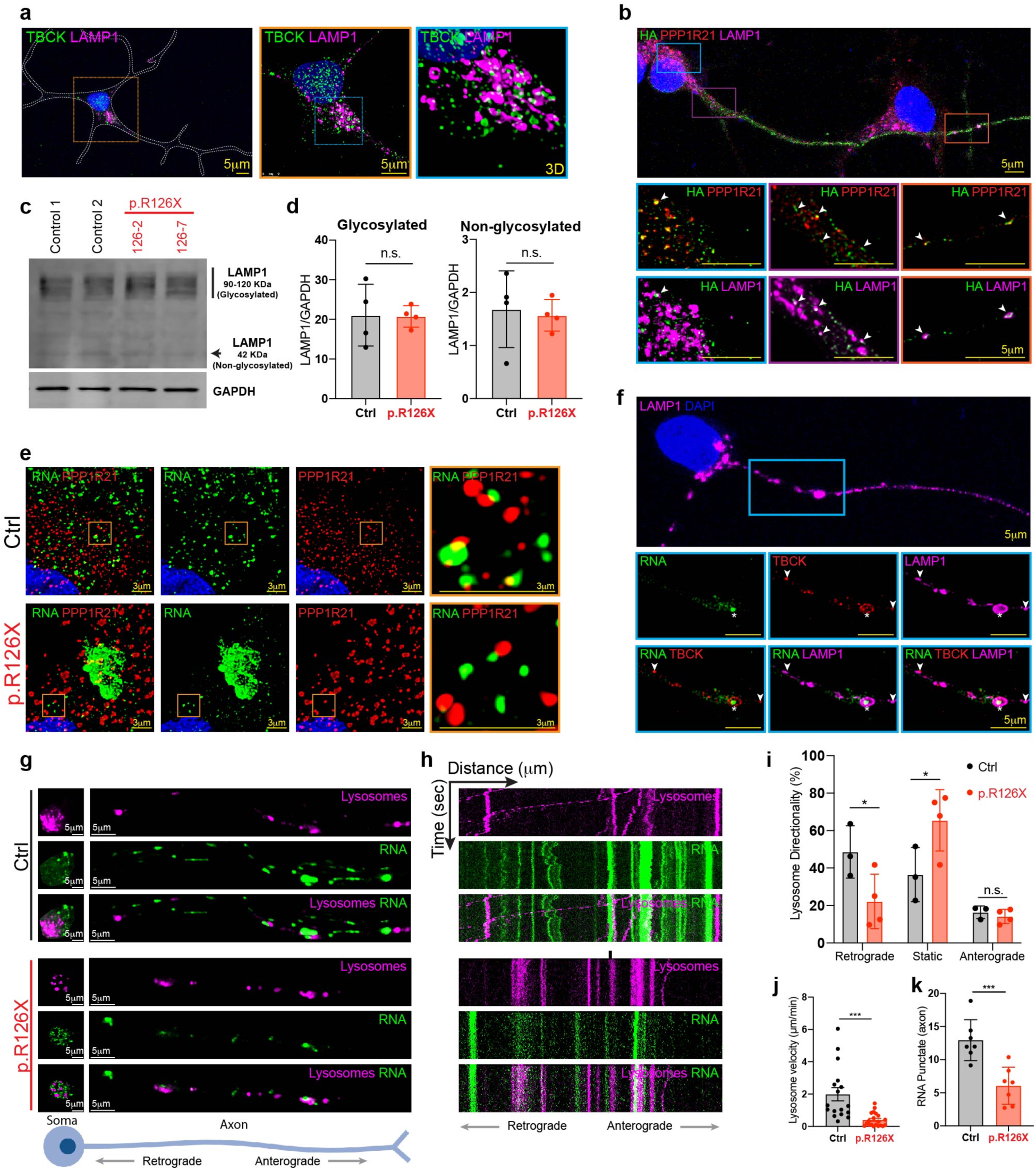
TBCK deficiency causes defects in lysosome formation and trafficking. **a,** Representative confocal images from a control iNeuron showing TBCK and LAMP1. Different inset magnifications (orange and blue squares) show the distribution and colocalization of TBCK and LAMP1 (3D= three-dimensional image). **b,** Confocal images of control neurons transduced with TBCK-HA and stained with anti-HA, PPP1R21 and LAMP1. Inset magnifications (blue, purple and orange) were taken from a long the neuron to show presence of FERRY complex (TBCK/PPP1R21) in lysosomes. Head arrows point colocalization of FERRY complex with lysosomes. **c**, Immunoblot of lysosomal marker (LAMP1) in control (Ctrl) and TBCK-deficient iNeurons (p.R126X). **d,** Quantification of LAMP1 (glycosylated and non-glycosylated) levels relative to GAPDH (n=4). Significance was calculated using unpaired t-test analysis. **e,** HyVolution confocal images in control and TBCK-deficient fibroblasts (p.R126X) showing PPP1R21 and RNA (Syto RNASelect). Different inset magnifications (orange squares) show the distribution and colocalization of PPP1R21 and RNA. **f,** Confocal images of control neurons labeled with Syto RNASelect to detect RNA and stained with anti-TBCK and LAMP1. Head arrows in the inset magnifications (blue squares) show colocalization of TBCK with LAMP1 and asterisks mark RNA within TBCK/lysosomes. **g,** Confocal images of control and TBCK-deficient neurons (p.R126X) plated on microfluid chamber showing lysosomes (LysoTracker) and RNA (Syto RNASelect) in soma and axons. **h,** Kymographs and analysis of (**i**) lysosome motility, (**j**) lysosome velocity and (**k**) RNA particles in control (Ctrl) and TBCK-deficient iNeurons (p.R126X) (n=3). Significance was calculated using unpaired t-test analysis. *p<0.01 and ***p<0.001. All graphs show error bars with mean ± SD from independent experiments (n).

### TBCK-deficient neurons have impaired lysosomal motility and mRNA trafficking

Endolysosomal vesicles, including late endosomes and lysosomes, have been shown to traffic mRNA, particularly in neurons^9^. Moreover, lysosomal trafficking defects are also linked to pediatric-onset neurodegeneration, reminiscent of TBCK deficiency ^29^. Hence, we hypothesized that TBCK deficiency may lead to mRNA trafficking and/or transport defects if TBCK indeed functionally disrupts the complex. To test this, we used the RNA-specific nucleic acid stain, sytoRNA in control and p.R126X fibroblasts. We found evidence that sytoRNA+ puncta appeared to cluster in the perinuclear area of p.R126X fibroblasts relative to controls, where the RNA signal is more dispersed along the cytoplasm (Fig. 7e). Moreover, sytoRNA+ puncta were not as clearly colocalized with PPP1R21 (i.e., early endosomes trafficking mRNA via the PPP1R21/FERRY complex) in p.R126X fibroblasts compared to controls (Fig. 7e).

In order to explore why neurons may be more susceptible to TBCK deficiency, we then asked whether disrupting the FERRY complex could have compartment-specific phentoypes. We cultured our iNeu in custom made microfluid chambers to clearly separate axons from the somatodendritic compartments^30^. We hypothesized that axonal compartments, where mRNA would need to travel greater distances from the nucleus, will be preferentially affected by mRNA transport defects. Indeed, we found that TBCK-deficient neurons have significantly fewer total sytoRNA+ puncta (RNA) specifically in axons, but not soma, when compared to control neurons (Fig. 7k).

To further probe if reduced axonal mRNA content was linked to impaired FERRY-mediated mRNA “hitchhiking” onto lysosomes, we performed live imaging to quantify trafficking of sytoRNA+ puncta and lysotracker+ vesicles in our human iNeu model (Fig. 7g). Kymograph analysis (Fig. 7h) revealed that axons in TBCK-deficient iNeu have a significantly higher percentage of static lysosomes (65%) relative to controls (36%), suggesting a lysosomal trafficking defect. Furthermore, axonal retrograde transport was significantly reduced in TBCK-deficient neurons, where only 22% of lysotracker+ puncta were moving in retrograde direction versus 49% in control axons (Fig. 7i). This could be consistent with disrupting JIP4 mediated dynein retrograde transport. Lysosome velocity in TBCK-deficient neurons is also significantly slower than controls (Fig. 7j). These experiments reveal that loss of TBCK leads to reduced axonal RNA content (Fig. 7k), likely by disrupting the assembly and/or trafficking of the FERRY complex along endolysosomal vesicles. Specifically, loss of TBCK leads to cell-type specific depletion of dynein motor adapter protein JIP4, and impaired lysosomal retrograde axonal trafficking

## DISCUSSION

Maintaining neuronal homeostasis and subcellular organization requires preserving functionally distinct cellular compartments. There is growing evidence that precise mRNA trafficking is key to delivering the proteins that sustain such localized function and communication between organelles. Recent evidence suggests that due to their unique morphology, neurons in particular utilize the machinery for intracellular organelle trafficking for mRNAs to “hitchhike” along organelles to be delivered for local protein translation^31^. There is evidence for mRNA trafficking along the surface of mitochondria^11^, as well as endolysosomal vesicles^8,9,32^. Defects in the endolysosomal pathway can manifest in numerous neurodegenerative diseases, both of pediatric and adult onset^29,33,34^. Here we revealed that disruption of the endolysosomal pathway, and specifically reduced axonal lysosomal and mRNA trafficking are hallmarks of TBCK Encephaloneuronopathy (TBCKE), a pediatric neurodegenerative disorder caused by TBCK deficiency (Fig. S7e).

TBCKE is caused by biallelic, usually loss of function variants, such as the Boricua founder variant c.376C>G (p.R126X) in exon 5 of TBCK. Typically, nonsense mutations lead to nonsense-mediated decay (NMD) of mRNA and hence loss of protein. However, we unexpectedly detected residual protein levels in patient derived neurons homozygous for the p.R126X variant. Nonsense-mediated mRNA decay efficiency can vary across transcripts^35,36^, depending on the position of the premature termination codon (PTC)^37,38^, tissue type^39^ or stress conditions^40,41^. A single-cell study revealed that this escape can be either by translational readthrough at the PTC or by a failure of mRNA degradation after successful translation termination at the PTC^42^. Additionally, TBCK mRNA is known to undergo alternative splicing, with 9 different transcripts reported. TBCK isoform C is encoded by the transcript variant 4 mRNA (NM_033115.5) and lacks exon 5^43^. This protein could be detected by the used antibody (NOVUS, NBP1-83166) that recognizes residues from L670 to S784 encoded by the exons 23-25. Recent long-read sequencing data in human brain suggests that >30 TBCK RNA isoforms are detectable, whose function and physiological relevance remain unclear^44^.

RNA transport is critical for neuronal homeostasis, since highly compartmentalized spatiotemporal protein expression is required^32,45^. The best-known mechanism described for cellular mRNA transport is via mRNA-binding proteins (mRBPs)^46,47^. Here, we describe the functional impact in neuronal mRNA transport of the recently discovered FERRY complex, which was reported to link early endosomes with mRNAs to traffic in the endosome surface^13,18^. Our data underscores that endolysosomal-dependent mRNA trafficking can be of particular relevance to neurons. Furthermore, we find evidence that TBCK closely associates to LAMP2+ vesicles in control neurons, suggesting the FERRY complex may not be restricted to early endosomes. We reveal that TBCK-deficiency disrupts trafficking in a compartment-specific fashion, with axonal mRNA and lysosomal content being significantly reduced in TBCK-deficient neurons while soma mRNA levels remain unchanged. Our data also suggests that TBCK is critical to for maintaining FERRY complex stability, as TBCK-deficiency leads to undetectable levels of another complex member, c12orf4. Therefore, our data strongly supports that reduced axonal mRNA trafficking, likely via disruption of the FERRY complex (due to TBCK and C12orf4 deficiency) is associated with severe, pediatric-onset neurodegeneration.

Previous data suggests that the FERRY complex is formed by five proteins. Three of them, PPP1R21, CRIZL1 and GATD1, were resolved by Cryo-EM. The remaining two, TBCK and C12ORF4, were detected by affinity chromatography^13,18^. We did not detect GATD1 in our neuron interactome data, and were not able to find reliable commercial antibodies to perform co-IP experiments. Therefore, we cannot conclusively prove or disprove the presence of GADT1 in our human neuronal model.

Our interactome data revealed novel interactions of TBCK protein with TRIM27 and JIP4. These proteins have been implicated in the retromer complex (TRIM27) and in lysosomal trafficking (JIP4 is a dynein motor adapter linked to lysosomes and autolysosomes)^48,49^. We also identified TBCK interactions with nuclear proteins such as MED23. It remains unclear at this point which of these interactor proteins could potentially associate with the FERRY complex. More likely, these data suggest that TBCK has additional roles, yet to be defined, beyond the FERRY complex, consistent with previous reports of links with mTORC1 pathway and autophagy, for instance. It is also conceivable that the complex may have additional functions beyond endolysosomal mRNA trafficking.

Our data also uncovers that TBCK deficiency is associated with enlarged early endosomal size and perinuclear distribution, a robust finding across human cell lines (neurons, fibroblasts and HEK293). Our findings suggest that TBCK could play a role in the maturation of endolysosomal vesicles, as we find increased content of vesicles expressing early endosomal markers (EEA1 and PPP1R21). Nevertheless total RAB5 levels are unchanged by immunoblot. This phenotype is reminiscent of RAB5 overactivation^50^. This raises the question of whether TBCK may regulate RAB5 activity, given that it remains unknown if the Rab Gap kinase domain of the protein is functionally active (versus being a pseudokinase). Interestingly, the interaction with TRIM27 may provide an alternative potential link to the enlarged early endosome phenotype. TRIM27, together with MAGE-L2, VPS35, Ube20 and USP7 inhibit WASH degradation and promote the endosomal actin nucleation by Arp2/3 complex^23,48^. Knockout of WASH produces aberrant vesicles positive for EEA1 and LAMP1, a similar phenotype observed in TBCK-deficient cells^51^. Further studies will help to elucidate how TBCK may interact with the retromer complex and if this could play a role in the endosomal phenotype here described. Remarkably, enlarged early endosomes have been identified as an early cytopathological marker of Alzheimer’s disease, further implicating endolysosomal dysfunction to neurodegeneration across the age spectrum^52^.

The role of mRNA transport defects as a potential common denominator across neurodegenerative disorders has been recently explored^32^. There is evidence for disrupted axonal mRNA transport across some of the common, typically sporadic and adult-onset disorders like ALS/FTD^9^ and Alzheimer’s Disease^53,54^. The literature largely implies disruptions in mRNP assembly or trafficking in these disorders^32,55,56^. However, there is less evidence regarding the physiological relevance of the more recently described endolysosomal-associated mRNA trafficking. Initially, endosomal mRNA trafficking was found as an alternative hitchhiking pathway for long-distance transport in Ustilago maydis^57^, but it seems to be a widespread mechanism along species^47,58^. In neurons, defects in endosomes and lysosomes lead to impaired mRNA transport, local protein synthesis, mitochondrial dysfunction, and loss of axon integrity^8,13,29^.

Ultra-rare monogenic disorders can provide a window to gain insight into the functional relevance of these novel trafficking complexes. In addition to TBCK, loss of function variants in PPP1R21 and C12orf4 are also linked to severe, pediatric-onset neurologic dysfunction. Although we hypothesize that it is likely that mRNA trafficking defects may also play a role in those disorders linked to the FERRY complex, further studies should test if axonal mRNA trafficking is similarly disrupted or if these proteins have other yet to be understood cellular functions. Although not directly involved in mRNA binding, recent evidence links the BORC complex to axonal mRNA trafficking defects and neurodegeneration by impairing axonal lysosomal transport ^29^. Similar to our data, they find mitochondrial defects, suggesting that the endolysosomal trafficking of mRNA may be particularly relevant to deliver nuclear-encoded mitochondrial gene products. Similar findings in the c9orf72 ALS models further supports that mRNA trafficking is crucial for axonal mitochondrial and overall neuronal homeostasis^59^. Therefore, our data further supports the concept that secondary mitochondrial dysfunction due to mRNA axonal trafficking defects may contribute to the selective neuronal vulnerability seen in neurodegenerative disorders. Further studies should address how organelle-mediated mRNA transport relates to other transport (RBPs, etc.) mechanisms, whether their functional relevance is potentially tissue or development specific, and what mediates the specificity of mRNA transcripts’ interaction with diverse cellular trafficking options.

In context of the FERRY complex, our work further expands the localization of FERRY beyond early endosomes and provides evidence for the first time that disrupting a member of the complex (TBCK) disrupts axonal lysosomal trafficking. Furthermore, our data suggests that FERRY may effectively link to the neuronal trafficking machinery via JIP4. Our data across different cell lines also suggests that the interaction of TBCK with JIP4 may be differentially regulated in neurons, as loss of TBCK does not lead to reduction of JIP4 levels in fibroblasts and HEK293 cells. This data further provides a potential mechanism for the selective neuronal vulnerability observed in TBCK deficiency. We find axonal lysosomal trafficking defects yet, to date, there is no evidence of JIP4 to be associated with early endolysosomal vesicles. JIP4 is reportedly involved in trafficking autolysosomes^49^ and lysosomes^60^ in neurons. Hence, we suspect the FERRY complex may be associated with endolysosomal compartments across maturation stages, from early endosome to mature lysosomes. On the other hand, JIP4 has been reported to be predominantly reported to mediate retrograde trafficking via its dynein interactions. Indeed, we observed TBCK-deficient human iPSC neuron axonal compartments had significant retrograde trafficking of lysotracker positive vesicles. Yet, we also observed reduced lysosomal content in the axon, suggesting an anterograde component of trafficking, or alternatively, a maturation defect, could also be playing a role. Further studies are needed to discern amongst these possibilities.

A limitation of our current study is that we were unable to demonstrate that restoring TBCK would rescue the described phenotypes. In particular, we wanted to test if the lysosomal trafficking defect could be rescued by restoring TBCK expression levels. We attempted rescuing the patient-derived iPSC lines, transducing with a lentiviral vector to reintroduce TBCK. Unfortunately, this approach was unsuccessful, resulting in extensive cell death in both patient-derived fibroblasts and iPSC cells. Furthermore, overexpression of TBCK in other cell types including primary fibroblasts, also led to cell death in both controls and mutant cells. Despite not having isogenic lines, we performed most of the phenotypic characterization across 3 different human cell lines and show a robust and consistent phenotype of endolysosomal defects in TBCK deficiency. Our experience suggests that overexpression of TBCK is not well tolerated and it is possible that the stoichiometry of the complex is tightly regulated in the cell. Similarly, transduction of JIP4 in our patient-derived neurons to try to rescue the motility phenotype was unsuccessful, as mutant cells did not survive transduction. TBCK may also have additional functions beyond the FERRY complex (as suggested by nuclear localization in neurons and our interactome data revealing nuclear and retromer complex-associated proteins) that become toxic with overexpression. These observations also have implications for future clinical therapeutic strategies for TBCK syndrome, as future gene replacement strategies may need to carefully consider expression levels and target cell types in the brain.

Overall, here we describe axonal-specific mRNA and lysosomal trafficking defects in patient-derived, human neuronal model of the pediatric neurodegenerative disorder TBCKE. We postulate that mRNA trafficking defects are due to disrupting the FERRY complex, as disease-causing variants result in not only TBCK deficiency, but we unexpectedly found depleted levels of c12orf4. Our data also reveals that TBCK directly interacts with motor adapter protein JIP4, and that TBCK-deficiency may regulate JIP4 levels in a tissue-specific fashion by depleting JIP4 levels in neurons but not fibroblasts. Hence our data provide novel insight into the pathophysiology of TBCKE, and provides further evidence that mRNA trafficking defects may contribute to neurodegeneration across the age spectrum.

## RESOURCE AVAILABILITY

### Lead Contact

Further information and requests for resources and reagents should be directed to and will be fulfilled by the Lead Contact, Xilma R Ortiz-González.

### Materials Availability

Stable human iPSC-derived neurons expressing Neurogenin-2 utilized in this study can be requested through Dr. Xilma R Ortiz-González upon Material Transfer Agreement.

### Data and Code Availability

All input data are freely available from public sources. Mass spectrometry proteomics data have been deposited to the ProteomeXchange Consortium via the PRIDE.

## EXPERIMENTAL MODEL AND SUBJECT DETAILS

### Human Induced Pluripotent Stem Cell lines

iPSCs were generated from patient fibroblasts homozygous for the Boricua mutation (p.R126X), obtained with the appropriate consent under an IRB-approved protocol. According to the clinical course of progressive encephalopathy and motor neuropathy, these patients were cataloged as severe^5^. Two individual lines from unrelated patients were used (126-2 and 126-7). Control cells (ICB4 and CHOP) are from healthy and unrelated to the affected individuals. iPSCs were transduced with lentivirus containing neurogenin-2 (Ngn2) at multiplicity of infection (MOI) 25 in presence of 5 μg/ml polybrene. This lentivirus (pLV[Exp]-TRE>mNeurog2[NM_009718.3](ns):T2A) was constructed and packaged by VectorBuilder (Vector ID: VB210325-1362twv). IPSCs were cultured and maintained in mTeSR (STEM-Cell Technologies, 85850).

### Generation of iPSC-derived neurons

Differentiation was driven by Ngn2 and according to the protocol previously described^20^. Briefly, iPSC-Ngn2 cells were dissociated and plated on Matrigel (BD Biosciences)-coated dishes with mTeSR containing Y-27632 (Tocris, 1254). On day 0, the culture medium was replaced with N2/DMEM-F12/NEAA (Invitrogen) containing human BDNF (10 mg/l, PeproTech), human NT-3 (10 mg/l, PeproTech), and mouse laminin (0.2 mg/l, Invitrogen, 23017015). Ngn2 was induced by doxycycline (2 μg/ml, Sigma, D3072) and maintained in the medium until day 10. On day1, puromycin (1 mg/l) was added to start the selection and maintained until day 2. On day 2, Neurobasal medium supplemented with B27/Glutamax (Invitrogen) containing BDNF and NT3, was added. On days 3-14, 50% of the medium was exchanged every other day. iPSC-derived neurons cells were used on day 14 in most experiments.

For stable expression of TBCK-HA in iPSC-derived neurons, a *TBCK-HA*-encoded lentivirus vector (pLV[Exp]-Neo-TRE>hTBCK[NM_001163435.3]/HA) was purchased from VectorBuilder (ID: VB221114-1285fwp). The lentivirus was transduced to iPSCs in presence of 5μg/ml polybrene for 36h and selected one week with mTeSR supplemented with 200 μg/ml G418. For the transgene expression, iPSCs were treated with 2 μg/ml doxycycline on day 1 during the differentiation to neurons.

### Primary patient fibroblasts

Fibroblasts were obtained with appropriate consent under an IRB-approved protocol at Children’s Hospital of Philadelphia. Cells were maintained in DMEM with glutamax media supplemented with 10-15% fetal bovine serum (FBS) and non-essential amino acids (NEAA). Experiments were carried out with cell of comparable passage number, and were not used past passage 18. Fibroblast line derived from patients homozygous for the Boricua mutation (p.R126X), with a clinical course of progressive encephalopathy and motor neuronopathy are defined as severe. Two lines from unrelated patients were used, 126-10 and 126-11. None of the p.R126X patients achieved language or independent ambulation. Control fibroblast cell lines (similar age individuals) were obtained from the NIGMS Human Genetic Cell Repository at the Coriell Institute for Medical Research.

### HEK293 cell culture

Cells were maintained in DMEM containing 10% FBS. The cells were maintained by passage into fresh media every 3–4 days. Cells were regularly tested for mycoplasma contamination using mycoplasma detection PCR (6601, Takara) and were negative for mycoplasma contamination. For stable expression of Flag-TBCK the 5’-3XFlag-TBCK-3’ fragment was excised from the Puc57 vector and gel-purified. The resulting fragment was subsequently ligated into the pQCXIN retroviral vector (Takara, catalog no. 631514). Retrovirus production was carried out in HEK293 cells (ATCC, VA). To generate TBCK knockdown lentiviral vectors (pLV[shRNA]-Neo-U6>hTBCK[shRNA) were constructed and packaged by VectorBuilder. Scramble shRNA, TBCK shRNA #1 (target sequence: CGGAATAGTGAAGACTTTATT; ID: VB221115-1377vmx), TBCK shRNA #2 (target sequence: GCTAAAGGCTTATCCATATAA; ID: VB221115-1378pxv) or TBCK shRNA #3 (target sequence: CCATCCCATCTCCTCAAATAT; ID: VB221115-1379pha) were transduced using Opti-MEM in presence of 5 μg/ml polybrene for 36h and selected with 200 μg/ml G418 in growing media for one week.

### Lymphoblastoid cell lines (LCLs)

Control and mutant (p.K213fs) LCLs were donated by Fowzan S. Alkuraya. Cell cultures were maintained in RPMI 1640 medium containing 20% FBS + 1% L-glutamine. Medium was replaced every 2-3 days to maintain the cell density between 1×10^5^-2×10^6^ cells/ml. Patient’s LCLs with C12orf4 mutation (p.K213fs), come from a study of 143 multiplex consanguineous families identified by whole-exome sequencing^16^.

## METHOD DETAILS

### Proteomic sample preparation

IPSC-derived neurons (control and p.R126X cells) were differentiated and grown in 150mm dishes. These were placed on ice, washed twice with cold PBS and harvested by scraping cells. Cells were centrifuged 12000 RPM for 10 min at 4 °C. Supernatant was removed and pellet was resuspended with lysis buffer (50 mM HEPES, pH 7.5, 150 mM NaCl, 0.5%Triton X-100, 1mM EDTA, 1mM EGTA, 10mM NaF, 2.5mM Na3VO4, complete protease/phosphatase inhibitor cocktail (Roche). Cell lysate was centrifuged 12,000 RMP for 10 min at 4 °C and supernatant was acetone precipitated overnight and stored at −20 °C. Protein pellets were solubilized, reduced and alkylated by addition of sodium deoxycholate buffer containing tris(2-carboxyethyl)phosphine and 2-chloroacetamide then heated to 95°C for 10 minutes. Proteins were then enzymatically hydrolyzed for 1.5 hours at 37 °C by addition of LysC and trypsin (Promega, Fujifilm Wako chemicals, USA). The resulting peptides were de-salted, dried by vacuum centrifugation, and reconstituted in 0.1% trifluoroacetic acid containing iRT peptides (Biognosys Schlieren, Switzerland) for LC MS/MS analysis.

### TBCK interactome assay

IPSC-derived neurons were lysed using RIPA buffer (Thermo Fisher Scientific, 89901) supplemented with protease/phosphatase inhibitor cocktail (Thermo Fisher Scientific, 1861281). Cells were centrifuged 12000 RPM for 10 min at 4 °C and supernatant was recovered. Protein concentration was determined by BCA method according to the manufacturer’s instructions. 1mg of protein was incubated with 1μg of anti-TBCK (Novus, NBP1-83166) or anti-IgG (Cell Signaling, 5415) overnight at 4°C in slow rotation. This mix was equilibrated to up 500 ul with immunoprecipitation (IP) buffer (225mM NaCl, 50mM Tris pH 8, 0.5% NP40 and 5mM EDTA). Next day, 25 μl of Protein A/G Magnetics Beads (Thermo Fisher Scientific, 88803) were added to the mix and incubated for 4h at 4°C in rotation. The beads were washed three times with ice-cold IP buffer and then proteins were eluted with 30 ul of 1x LDS buffer. Proteins were heated at 95°C for 5 min and place on ice for other 5min. Samples were loaded onto NuPAGE 12% (Thermo Fisher Scientific) and run for 15 min. The gels were stained with Coomassie Blue and destained with acetic acid solution (10%). Each gel line was excised, reduced with TCEP, alkylated with iodoacetamide, and digested with trypsin. Trypsin digestion were analyzed by LC MS/MS.

### LC-MS/MS analysis

Proteome and interactome were analyzed by LC/MS/MS using a QExactive HF mass spectrometer coupled with an Ultimate 3000 nano ultra-performance liquid chromatography system and an EasySpray source (Thermo Fisher Scientific; San Jose, CA). Peptides were loaded onto a 75 µm x 2 cm trap column (Acclaim PepMap 100; Thermofisher) at 5 μl/min, and separated by reverse phase high performance liquid chromatography on a 75 μm internal diameter × 50cm 2µm PepMap rapid separation liquid chromatography C18 column (Thermo Fisher Scientific). Mobile phase A consisted of 0.1% formic acid and mobile phase B of 0.1% formic acid/acetonitrile. The gradient was started at 1% phase B from 0-3 minutes, 5% phase B at 5 minutes, 15% B at 15 minutes, 45% at 155 minutes before increasing to 99% B to wash column off. Flow rate started at 300 nl/min, was lowered to 210 nl/min from 15.1 to 155 minutes and increased back to 300 nl/min. Data was acquired using data independent acquisition (DIA). Mass spectrometer settings were: one full MS scan at 120,000 resolution and a scan range of 300-1650 m/z with an automatic gain control (AGC) target of 3×10^6^ and a maximum inject time of 60 ms. This was followed by 22 (DIA) isolation windows with varying sizes at 30,000 resolution, an AGC target of 3×10^6^, and injection times set to auto. The default charge state was 4, the first mass was fixed at 200 m/z and the normalized collision energy for each window was stepped at 25.5, 27 and 30. The suitability of Q Exactive HF instrument was monitored using QuiC software (Biognosys; Schlieren, Switzerland) for the analysis of the spiked-in iRT peptides. As a measure for quality control, we injected standard E. coli protein digest in between samples (one injection after every four biological samples) and collected the data in data dependent acquisition (DDA) mode. The collected DDA data were analyzed in MaxQuant and the output was subsequently visualized using the PTXQC package to track the quality of the instrumentation. Statistical analysis of acquired data were performed as described below. The raw files for DIA analysis were processed with Spectronaut version 15.1 in DirectDIA mode using the Prot human proteome database (06/04/2021). The default settings in Spectronaut were used for peptide and protein quantification. Perseus (1.6.14.0) was employed for data processing and statistical analysis using the MS2 intensity values generated by Spectronaut. The data were log2 transformed and normalized by subtracting the median for each sample. Student’s t-test with was employed to identify differentially abundant proteins using p value < 0.05 as significant threshold.

### Functional enrichment analysis

Gene ontology (GO) term enrichment analysis was input into ShinyGO v0.77 (http://bioinformatics.sdstate.edu/go/) with human genome as the background. False discovery rate (FDR) values were calculated using the Benjamini–Hochberg test. P values were calculated using Fisher’s exact test in ENRICHR. Selected and significantly enriched (P < 0.05 (-log10=1.301) with Benjamini–Hochberg correction). GO annotations for biological processes are represented as bar graphs. Graphs include terms in all categories (biological processes, molecular function and cellular component). Due to the hierarchical nature of GO terms in ShinyGO (i.e. groups of terms have a nested nature to assign relationships between them) we only considered the most proximal term in each hierarchy to ensure terms were specific and directly comparable. Terms were ranked by FDR value and non-redundant top terms were included in each figure.

### Mitochondrial respiration assay

20,000 iPSC-derived neurons per well were seeded into Matrigel coated 96 well Seahorse cell culture plates (Agilent) and assayed at day 14 of plating. Prior to the assay, plates were switched to a 37C non-CO2 incubator for 1 hour, in Seahorse experiential medium (Seahorse Basal DMEM medium with 25 mM glucose, 2 mM L-glutamine, and 10 mM sodium pyruvate, ph7.4 at 37C). Seahorse Basal DMEM medium was used due to its bicarbonate free, low in buffer capacity, and low phenol red content. Via XF-96 Extracellular Flux Analyzer (Agilent); Oxygen consumption rates (OCR) and extracellular acidification rates (ECAR) were measured under a sequential drug treatment involving Oligomycin A (1.25 μM, an inhibitor of ATPase), Carbonyl cyanide 4-(trifluoromethoxy) phenylhydrazone (FCCP, 1 μM, a mitochondrial uncoupler), and a combination of Rotenone (1 μM, an inhibitor of mitochondrial complex I) and Antimycin A (1.8 μM, an inhibitor of mitochondrial complex III). Reads were normalized to protein content.

### Immunoprecipitation assay

Cells were lysed using RIPA buffer (Thermo Fisher Scientific, 89901) supplemented with protease/phosphatase inhibitor cocktail. Cells were centrifuged 12000 RPM for 10 min at 4 °C and supernatant was recovered. Protein concentration was determined by BCA (Thermo Fisher Scientific, 23225) method according to the manufacturer’s instructions. 500 μg of protein was incubated with 1μg of anti-TBCK (Novus, NBP1-83166) anti-PPP1R21 (Atlas Ab, HPA036791), anti-Flag (Sigma, F1804), anti-HA (Thermo Fisher Scientific, 26183) or anti-IgG (Cell Signaling, 5415) overnight at 4°C in slow rotation. This mix was equilibrated to up 500 ul with immunoprecipitation (IP) buffer (225mM NaCl, 50mM Tris pH 8, 0.5% NP40 and 5mM EDTA). Next day, 25 μl of Protein A/G Magnetics Beads (Thermo Fisher Scientific, 88803) were added to the mix and incubated for 4h at 4°C in rotation. The beads were washed three times with ice-cold IP buffer and then proteins were eluted with 30 μl of 1x LDS buffer (Invitrogen NP0007). Proteins were heated at 95°C for 5 min and place on ice for other 5min.

### Immunoblotting

Protein concentration was determined by BCA (Thermo Fisher Scientific, 23225) method according to the manufacturer’s instructions. Protein samples were diluted in an equal volume of 4x LDS sample buffer (Invitrogen, NP0007) and supplemented with DTT to a final concentration of 50mM (Bio-Rad, 1610611). Protein samples (30 μg) were separated on NuPAGE 4-20% Tris-Glycine gels (Thermo Fisher Scientific), transferred to PVDF membranes (Bio-Rad, 1620177) and blocked with 5% milk in TBS + 0.1% Tween (TBS-T) for 2h. After washing 3 times with TBS-T the blots were incubated with primary antibodies overnight at 4°C. Blots were washed 3 times with TBS-T and incubated with secondary antibodies (LI-COR Biosciences) for 2 hours at RT, washed again for 3 times in TBS-T, followed by LI-COR system detection, bands were quantified using Image Studio (LI-COR Biosciences). The following antibodies were used for western blot analysis: rabbit anti-LC3B (1:1000, Cell signaling, 27755), rabbit anti-p62 (1:1000, Cell signaling, 51145), mouse anti-mTOR (1:1000, Cell signaling, 45175), rabbit anti-phospho mTOR (1:1000, Cell signaling, 55365), rabbit anti-RAB5 (1:1000, Cell signaling, 35475), rabbit anti-Rab7 (1:1000, Cell signaling, 9367T), rabbit anti-TUJ1 (1:1000, Cell signaling, 5568), rabbit anti-Tom20 (1:1000, Cell signaling, 72610), mouse anti-Flag (1:1000, Sigma, F1804), goat anti-GAPDH (1:2000, RD Systems, AF5718), rabbit anti-TBCK (1:1000, Novus, NBP1-83166), rabbit anti-VDAC (1:1000, Abcam, ab15895), mouse anti-LAMP1 (1:500, DSHB, H4A3), rabbit anti-Cyt B (1:1000, Proteintech, 55090-1-AP), rabbit anti-RAB7 (1:1000, Cell signaling, 9367T), rabbit anti-PPP1R21 (1:1000, Atlas Ab, HPA036791), rabbit anti-C12ORF4 (1:1000, Sigma, HPA037871), rabbit anti-JIP4 (1:1000, Cell signaling, 5519), rabbit anti-TRIM27 (1:1000, IBL, 18791), rabbit anti-CRYZL1 (1:500, Novus, NBP1-89367), OXPHOS cocktail (1:500, Abcam, ab110413), goat anti-rabbit IgG (1:3000, LI-COR, 926-32211), goat anti-mouse IgG (1:3000, LI-COR, 926-68070), donkey anti-goat IgG (1:3000, LI-COR, 926-68074)

### Immunocytochemistry

Cells on coverslips were washed with PBS and fixed with 4% paraformaldehyde (PFA) for 10 mins at room temperature. PFA was removed and cells were washed 3 times with PBS + 0.1% Tween (PBS-T). Cells were incubated with blocking solution (0.1% Tween-20, 2% bovine serum albumin in PBS) for 1 hour, room temperature. The blocking solution was removed and replaced with primary antibodies prepared in blocking solution. After 3 washes with PBS-T, cells were then incubated with secondary antibodies in blocking solution, for 1 hour at room temperature, followed by PBS-T washes, 3 times. The following antibodies were used: rabbit anti-LC3B (1:5000, Cell signaling, 27755), mouse anti-p62 (1:500, Abcam, ab56416), mouse anti-RAB5 (1:500, Cell signaling, 2157), chicken anti-TUJ1 (1:1000, Aves labs), mouse anti-Flag (1:1000, Sigma, F1804), rabbit anti-TBCK (1:1000, Novus, NBP1-83166), mouse anti-LAMP1 (1:250, DSHB, H4A3), goat anti-LAMP2 (1:250, R&D Systems, AF6228), rabbit anti-PPP1R21 (1:500, Atlas Ab, HPA036791), rabbit anti-C12orf4 (1:500, Sigma, HPA037871), rabbit anti-JIP4 (1:500, Cell signaling, 5519), rabbit anti-TRIM27 (1:500, IBL, 18791), mouse anti-EEA1 (1:500, BD Tranduction), goat anti-mouse IgG Alexa Fluor 488 (1:500, ThermoFisher, A-11001), goat anti-mouse IgG Alexa Fluor 555 (1:500, ThermoFisher, A-21422), goat anti-mouse IgG Alexa Fluor 647 (1:500, ThermoFisher, A-21235), goat anti-rabbit IgG Alexa Fluor 488 (1:500, ThermoFisher, A-11008), goat anti-rabbit IgG Alexa Fluor 555 (1:500, ThermoFisher, A-21428), goat anti-rabbit IgG Alexa Fluor 647 (1:500, ThermoFisher, A-21245), donkey anti-goat IgG Alexa Fluor 488 (1:500, ThermoFisher, A-11055), donkey anti-goat IgG Alexa Fluor 555 (1:500, ThermoFisher, A-21432), donkey anti-goat IgG Alexa Fluor 647 (1:500, ThermoFisher, A-21447), goat anti-chicken IgG Alexa Fluor 488 (1:500, ThermoFisher, A-11039), goat anti-chicken IgG Alexa Fluor 555 (1:500,

ThermoFisher, A-21437), goat anti-chicken IgG Alexa Fluor 647 (1:500, ThermoFisher, A-21449).

### Quantitative PCR (qPCR)

RNA was isolated from eluted samples using the RNeasy Mini kit (QIAGEN, 74101). cDNA was prepared from total RNA using random hexamers and the SuperScript III First-Strand Synthesis system (ThermoFisher, 18080051). Relative gene expression of the cDNA was assayed by qPCR (software) using SYBR green PCR master mix (ThermoFisher, 4367659) following the manufacturer’s recommendations. Cycle counts for mRNA quantification were normalized to Gapdh. Relative expression (ΔCT) and quantification (RQ = 2 − ΔΔCT) for each mRNA were calculated using the ΔΔCT method as suggested. The following primers were used for qPCR: *gria1*, 5’-CTGAAGTGTGGGGATTATTTCCA-3’ and 5’- CTCAAAGCTGTCGCTGATGT-3’; *gria2*, 5’-TACAGATAGGGGGGCTATTTCC-3’ and 5’-GTTTGCCACCTCCAAATTGTC-3’; *gria3*, 5’-CCATCAGCATAGGTGGACTTT-3’ and 5’-TGGTGTTCTGGTTGGTGTT-3’; *grin4*, 5’-CGCTCCGCGCCATCGTC-3’ and 5’-CGCGCGCTCTCTCTCTTTC-3’; *grin1*, 5’- TCTCCAGCCAGGTCTACG-3’ and 5’-CACGGGTATGCGGTAGAAG-3’; *grin2a*, 5’- GGGCTGGGACATGCAGAAT-3’; *grin2b*, 5’-TTCCGTAATGCTCAACATCATGG-3’ and 5’-TGCTGCGGATCTTGTTTACAAA-3’; *grin2c*, 5’-GCTGGAAGAGCGGCCCTTTGT-3’ and 5’-CGCTGCTGAAGGTGTGGTTGCTCT-3’; *grin2d*, 5’- CTGCAGCCAGTGGACGACACG-3’ and 5’-GGGTTCGGTTGAGCTGGCTCCG-3’; *grin3a*, 5’-GCCACTCCACTGGACAATGTGGC-3’ and 5’- TTCGCCCCTTGGGAGTCAAACCA-3’; *tbck,* 5’- GTTGGACCG AAAGGGACATA-3’ and 5’-CCCTATTGGGAAATCAACATCATC-3’; *c12orf4,*5’-

GCTCCCTTAGTGAGGAAATTAAAGT-3’ and 5’- CATAAATCCTTCCAATTGCTTCTACAG-3’; *jip4,* 5’-GCTGACCAGATTAGCAGACTT-3’ and 5’-CTCAGTGTGTCTTTGATGTAATGC-3’; *trim27,* 5’- GGGCTTCAAGGAGCAAATC-3’ and 5’-GCCCGACGTCTCTTCTTTA-3’; *gapdh,* 5’- ACA TCGCTC AGACAC CATG-3’ and 5’- TGTAGTTGAGGTCAATGAAGGG-3’.

### Microfluid device preparation and culture

The Micro Fluidic Chambers wafers were designed and generously shared by the Song lab^61^. They were used to create PDMS microfluidic devices. SYLGARD 184 Silicone elastomer base mixed (Dow, 04019862) was used with a curing agent at a ratio of 9 to 1 grams of components A and B. The components were mixed by hand and then poured into the micro fluidic mold. Then the mold with the mixture was placed inside a Bel Art vacuum desiccator for at least 1 hour to remove air bubbles from the chambers. Then wafer with PDMS was placed on a hot plate at 80⁰C for 1 hour to cure. On the next day each individual chamber was cut and extracted from the wafer and cut to the appropriate size. Media reservoirs were punched into the chamber using a Biopsy punch with a diameter of 8 mm (Integra, 33-37). They were then given at least two cycles of UV before being used. Coat the glass bottom petri dishes (Celvis, D60-30-1.5-N) with PLO solution and incubate at 37⁰C overnight. The PLO was removes, dried to insert chambers. The chambers were then coated with Laminin overnight. Seed cells at a confluency of 5,000-10,000 cells, by dispensing the cells directly into the channel that connects the soma chamber to the media reservoir. Once neurites have already grown you can create a difference in volume between the two soma reservoirs and the two axonal reservoirs to promote growth through the chambers.

### Live-neuron imaging and motility analysis

For live-cell imaging, neurons were rinsed three times and transferred to low fluorescence Hibernate medium supplemented with 2% B27 and 0.5 mM GlutaMAX. Cells were visualized with a 40x/1.3 NA oil immersion objective on a Zeiss LSM880 confocal microscope. Time-lapse imaging was performed in an incubation chamber maintained at 37°C. To quantify motility, kymographs were generated using ImageJ as previously described^62^. Organelles were considered stationary if the net displacement was ≤ 10 μm during the entire acquisition period. If the net displacement was ≥ 10 μm throughout this period, the organelle was considered motile in its corresponding direction.

### Quantification and statistical analysis

All experimental data shown in this report, including immunofluorescence micrographs, were analyzed from at least three independent experiments or at least eight cells. The n number for each experiment, details of statistical analysis and software are described in the figure legends or main text. Statistical analyses used in this study include Student’s t-test and Fisher’s exact test. Statistical significance is defined as, n.s., not significant, *p < 0.05, **p < 0.01, ***p < 0.001. Statistical analysis was performed using Prism (GraphPad).

## Acknowledgments

We are grateful to the TBCK families that generously contributed samples for the generation of cell lines. The C12orf4 cell lines were generously shared by Dr. Fowzan Akuraya. Microfluidic chambers were designed and provided by the Song lab (supported by the NIH grant (1R01NS126541) and the Pennsylvania Department of Health grant (4100088540) to Y.S.). XOG is supported by NIH (K02NS112456, R01NS132795), the Burroughs Wellcome fund (CAMS award) and the TBCK foundation via the Orphan Disease Center at the University of Pennsylvania (Million Dollar Bike Ride Award to XOG). We are also grateful for the Center for Mitochondrial and Epigenomic Medicine and the Center for Molecular Therapeutics for access to shared equipment and imaging resources.

## Author Contributions

Conceptualization, XOG and MFM; Methodology, JATH, LM, TYL, YS; Investigation, MFM, JATH, LRR, Resources, FSA, YS; Writing-Original Draft, MFM, XOG, JATH; Writing-Review and Editing, XOG, YS, XOG,; Visualization, MFM, LRR, JATH; Funding Acquisition, Supervision and Project Administration, XOG.

## Competing Interests

Nothing to disclose

## Supplementary Figures

**Fig. S1:**
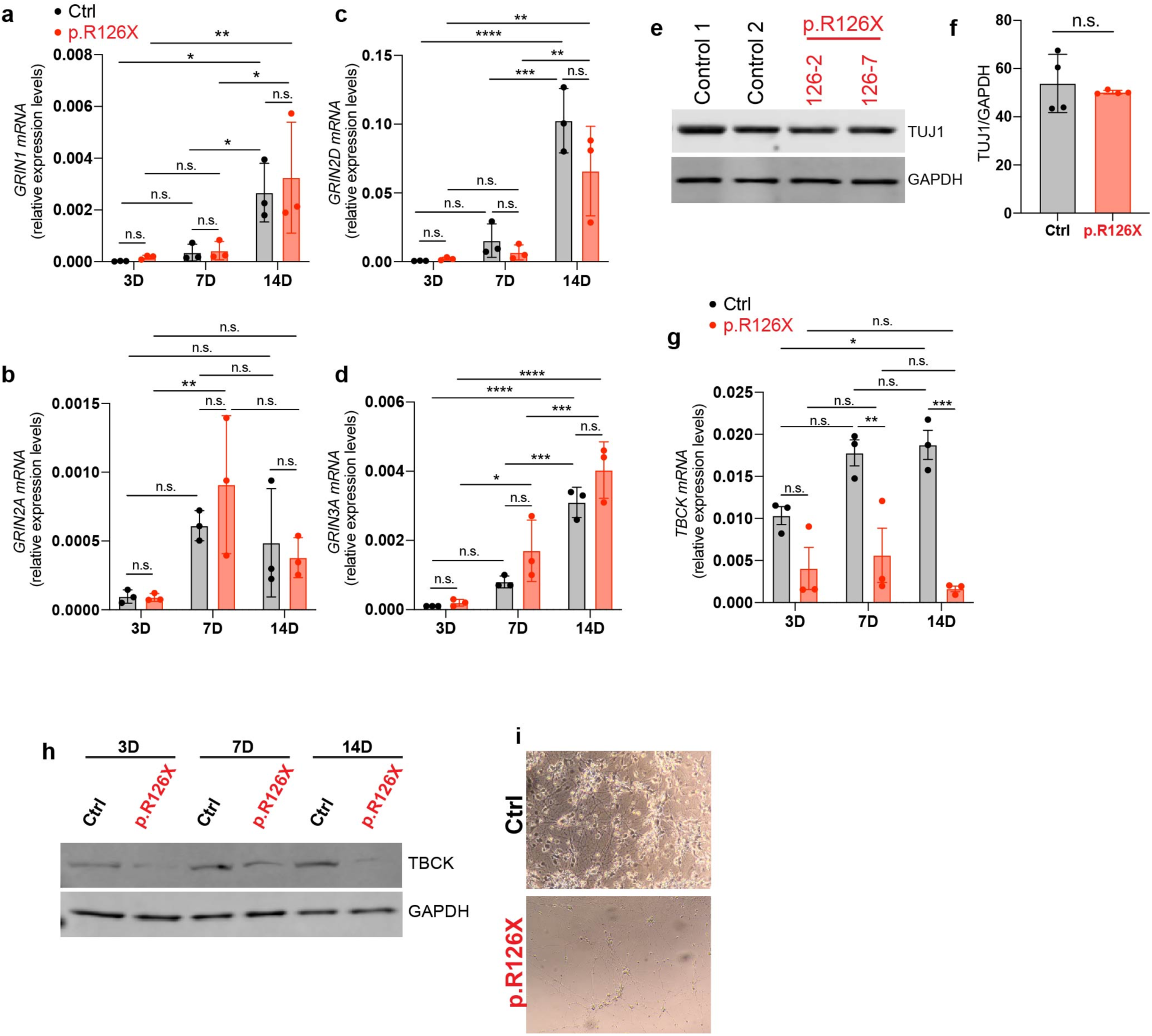
iPS cells show neuronal features after their differentiation using the NGN2 system. **a-d,** mRNA expression of glutamatergic receptors (*GRIN1, GRIN2A, GRIN2D AND GRIN3A*) at different stage (3,7 and 14 days) during the iPSC-derived neuron (iN) differentiation (n=3). Significance was calculated using two-way ANOVA with Sidak’s post hoc analysis for multiple comparison. **e,f,** Immunoblot of neuronal marker TUJ1 in iN at 14 days of differentiation (14D) in control (Ctrl) and TBCK-deficient neurons (p.R126X) (n=4). Significance was calculated using unpaired t-test analysis. **g,** mRNA expression of *TBCK* in iN at different stages during the differentiation in control (Ctrl) and TBCK-deficient neurons (p.R126X) (n=3). Significance was calculated using two-way ANOVA with Sidak’s post hoc analysis for multiple comparison. **h,** Immunoblot of TBCK during the differentiation of iN (3, 7 and 14 days). **i,** Micrography of iN culture during differentiation at stage of 14 days in vitro. *p<0.01, **p<0.05, ***p<0.001 and ****p<0.001. All graphs show error bars with mean ± SD from independent experiments (n).

**Fig. S2:**
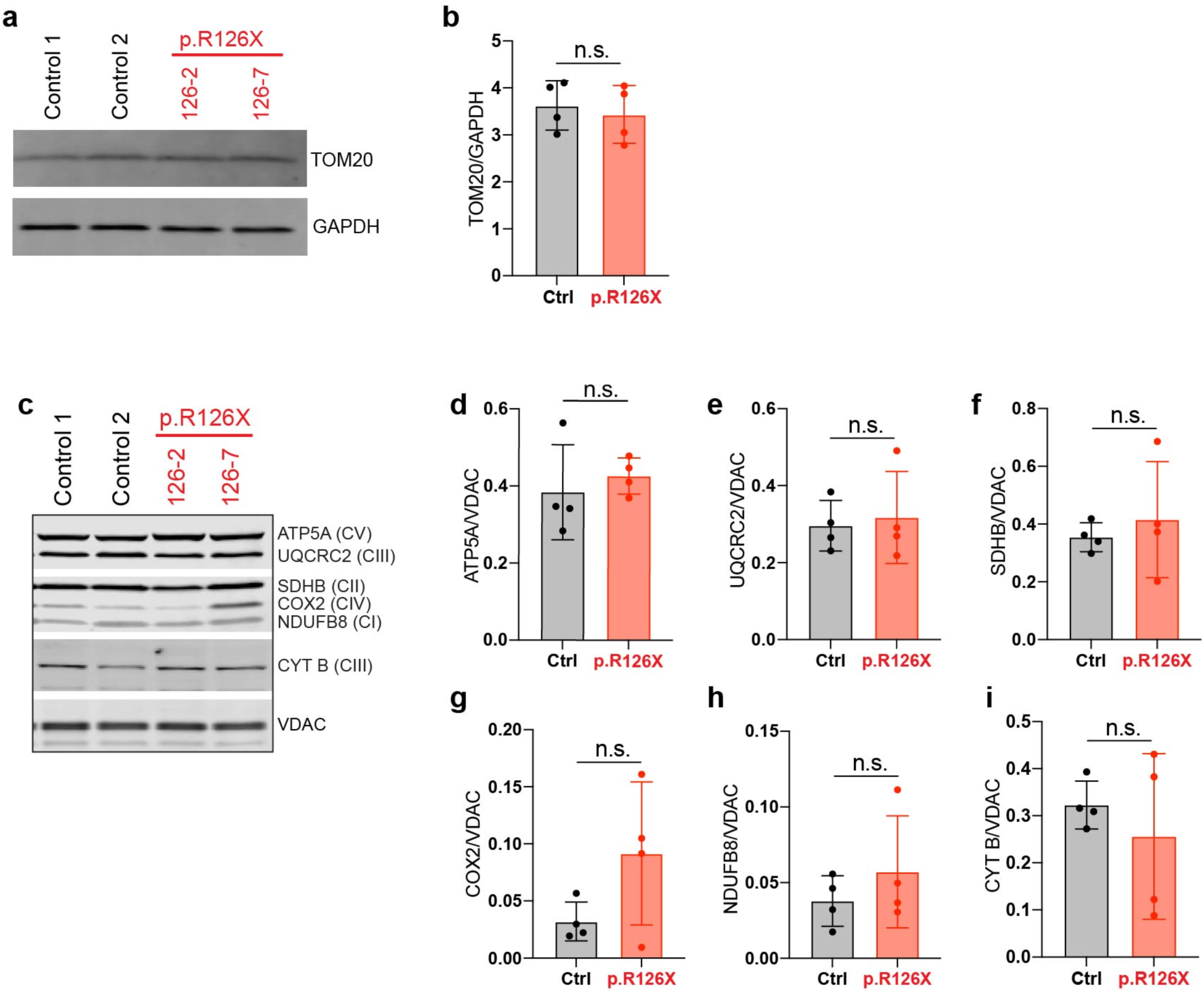
p.R126X mutation does not alter levels of mitochondrial proteins in neurons. **a,** Immunoblot of mitochondrial marker, TOM20, in control (Ctrl) and TBCK-deficient neurons (p.R126X) at 14D. **b,** Quantification of TOM20 relative to GAPDH (n=4). **c,** Immunoblot of proteins from the oxidative phosphorylation complexes (I-V) in iNeurons at 14D in control (Ctrl) and TBCK-deficient neurons (p.R126X) (n=4). Significance was calculated using unpaired t-test analysis. **d-i,** Quantification of immunoblot in **a** relative to VDAC (n=4). Significance was calculated using unpaired t-test analysis. All graphs show error bars with mean ± SD from independent experiments (n).

**Fig. S3:**
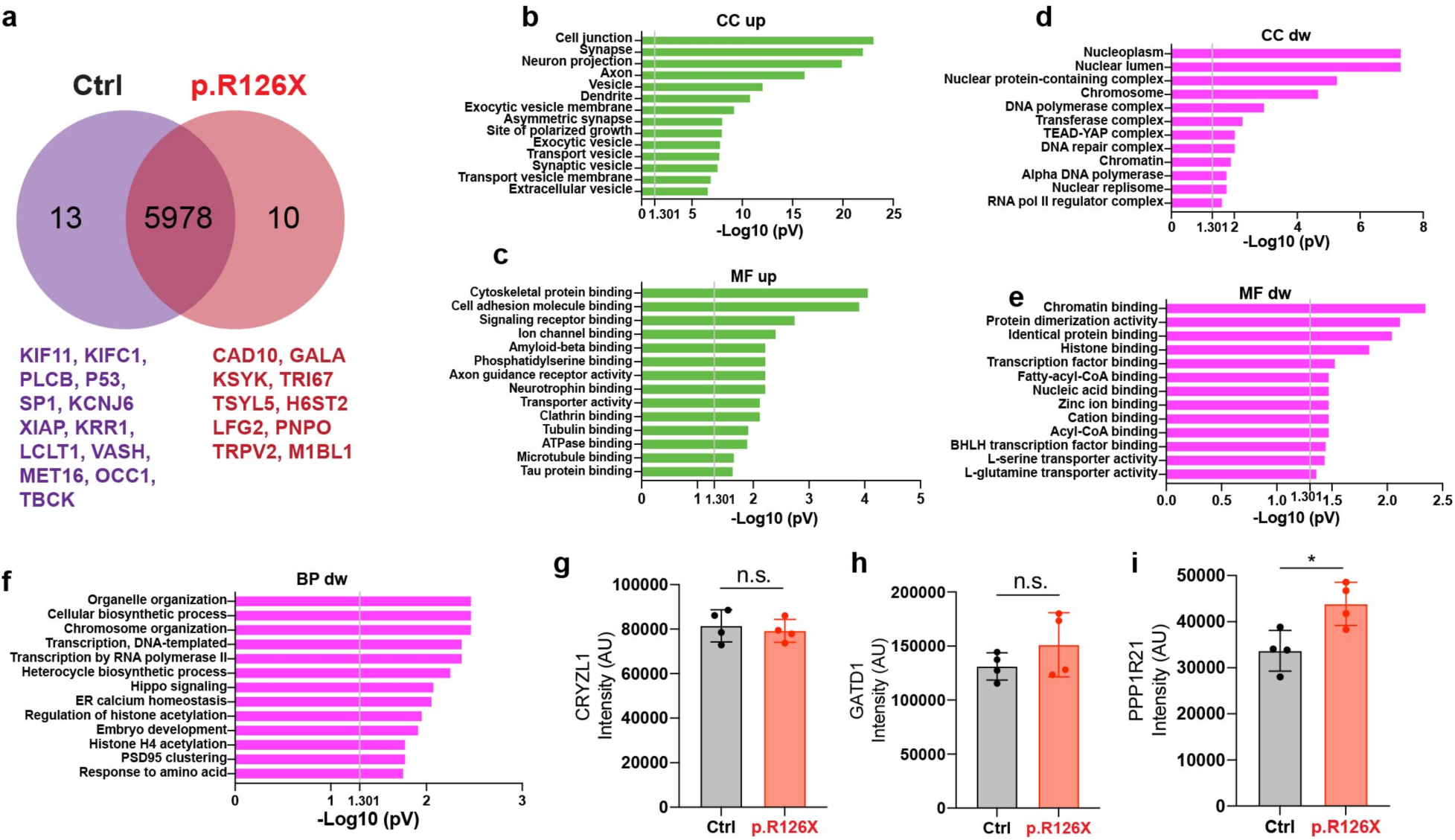
Protein levels obtained by immunoblots correlate with proteomics data. **a,** Venn diagram of proteomics between control and TBCK-deficient neurons (p.R126X). **b,c,** GO classification based on cellular component (CC) and molecular function (MF) of upregulated proteins from proteomics data. **d-f,** Classification of downregulated proteins from proteomics data based on cellular component (CC), molecular function (MF) and biological process (BP). **g-i,** Differential expression between control and TBCK-deficient neurons (p.R126X) obtained from the proteomics data (AU=arbitrary units, n=4). Significance was calculated using unpaired t-test analysis. Significantly enriched GO terms are shown with Benjamini-Hochberg FDR-corrected p-values. *p<0.01. All graphs show error bars with mean ± SD from independent experiments (n).

**Fig. S4:**
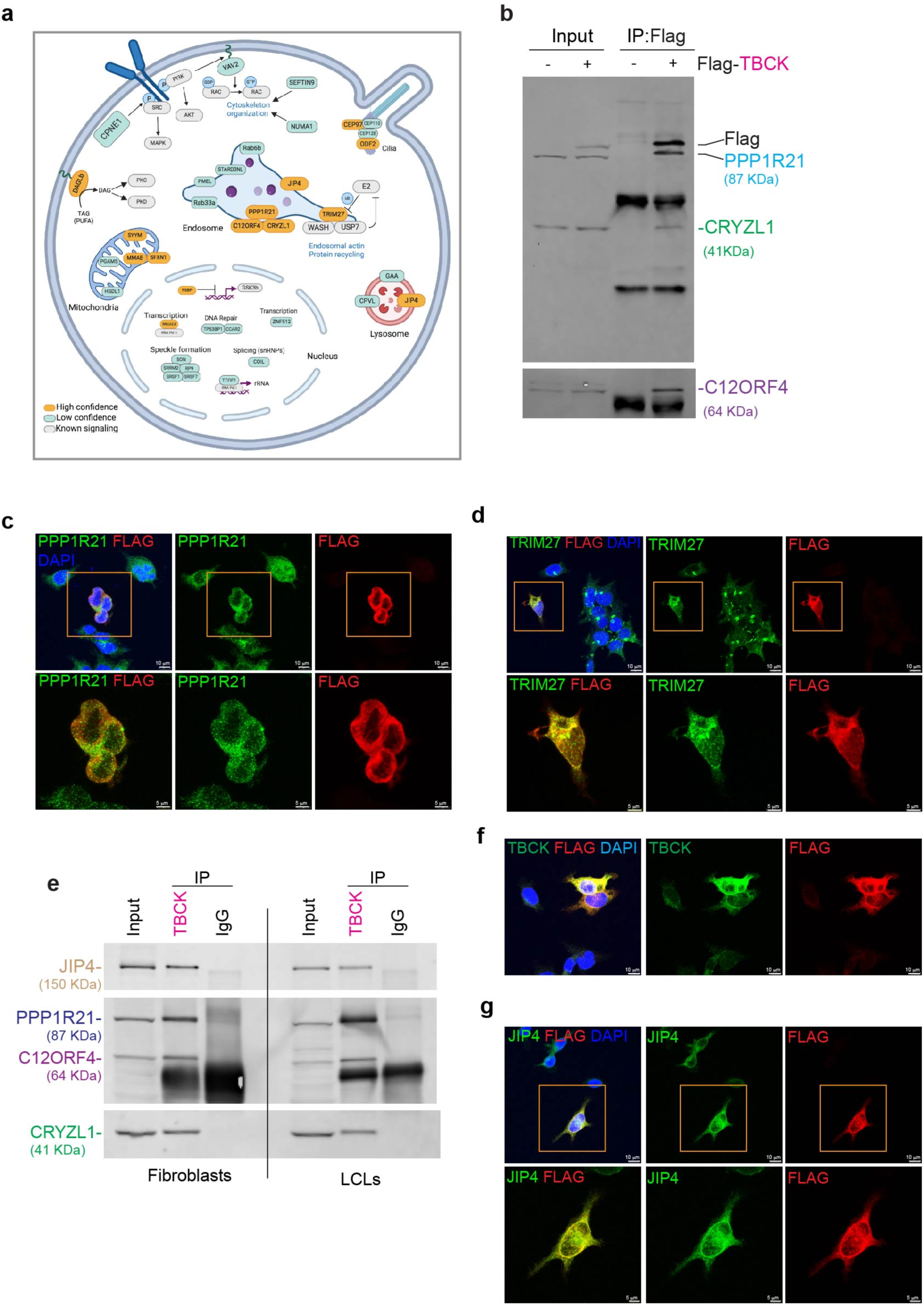
TBCK interacts with proteins associated with the endolysosomal pathway. **a,** Schematic representation of TBCK interactors in different cell compartments. Proteins in orange color were classified interactors with high confidence and in blue interactors with low confidence based on scores described in the text. Proteins in gray color are pathways and proteins described in other papers associated with the interactors. **b,** Immunoblot of proteins pulled down with Flag in HEK cells transduced (+) or not (-) with Flag-TBCK. Inputs were loaded with 20μg of total proteins. **c,** Representative immunostaining of Flag with PPP1R21 and (**d**) TRIM27 in HEK cells transduced with Flag-TBCK. Inset magnifications are label in orange squares. **e,** Representative immunoblot of TBCK immunoprecipitated from fibroblasts (left panel) and Lymphoblastoid cell lines (LCLs) (right panel). Inputs were loaded with 20μg of total proteins. **f,** Representative immunostaining of Flag with TBCK and (**g**) JIP4 in HEK cells transduced with Flag-TBCK. Inset magnifications are label in orange squares.

**Figure S5.**
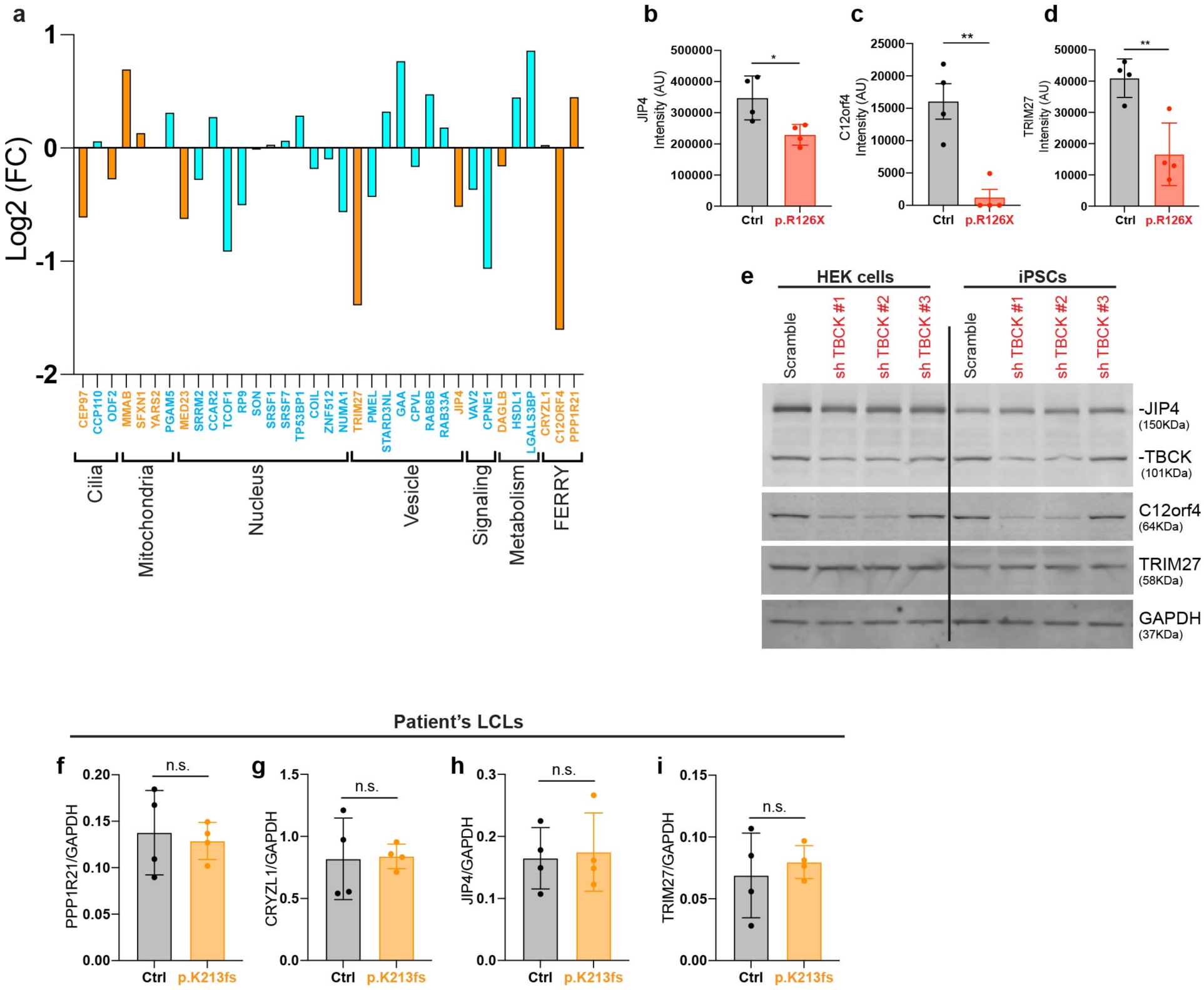
TBCK regulates C12ORF4 across different cell types. **a,** Abundance of TBCK interactors obtained by mass spectrometry of TBCK-deficient neurons (p.R126X). **b,** Differential expression of JIP4, (**c**) C12orf4 and (**d**) TRIM27 between control (Ctrl) TBCK-deficient neurons (p.R126X) obtained from the proteomics data. AU=arbitrary units (n=4). Significance was calculated using unpaired t-test analysis. **e,** Immunoblot of TBCK knockdown cells. HEK293 cells (left panel) and iPSCs (right panel) transduced with small hairpin RNA(shRNA) Scramble and three ShRNAs targeting TBCK (ShTBCK#1, ShTBCK#2 and ShTBCK#3). **f,** Quantification of protein levels of PPP1R21, (**g**) CRYZL1, (**h**) JIP4 and (**i**) TRIM27 in control (Ctrl) and affected LCLs (p.K213fs). Significance was calculated using unpaired t-test analysis. *p<0.01 and **p<0.05. All graphs show error bars with mean ± SD from independent experiments (n).

**Fig. S6:**
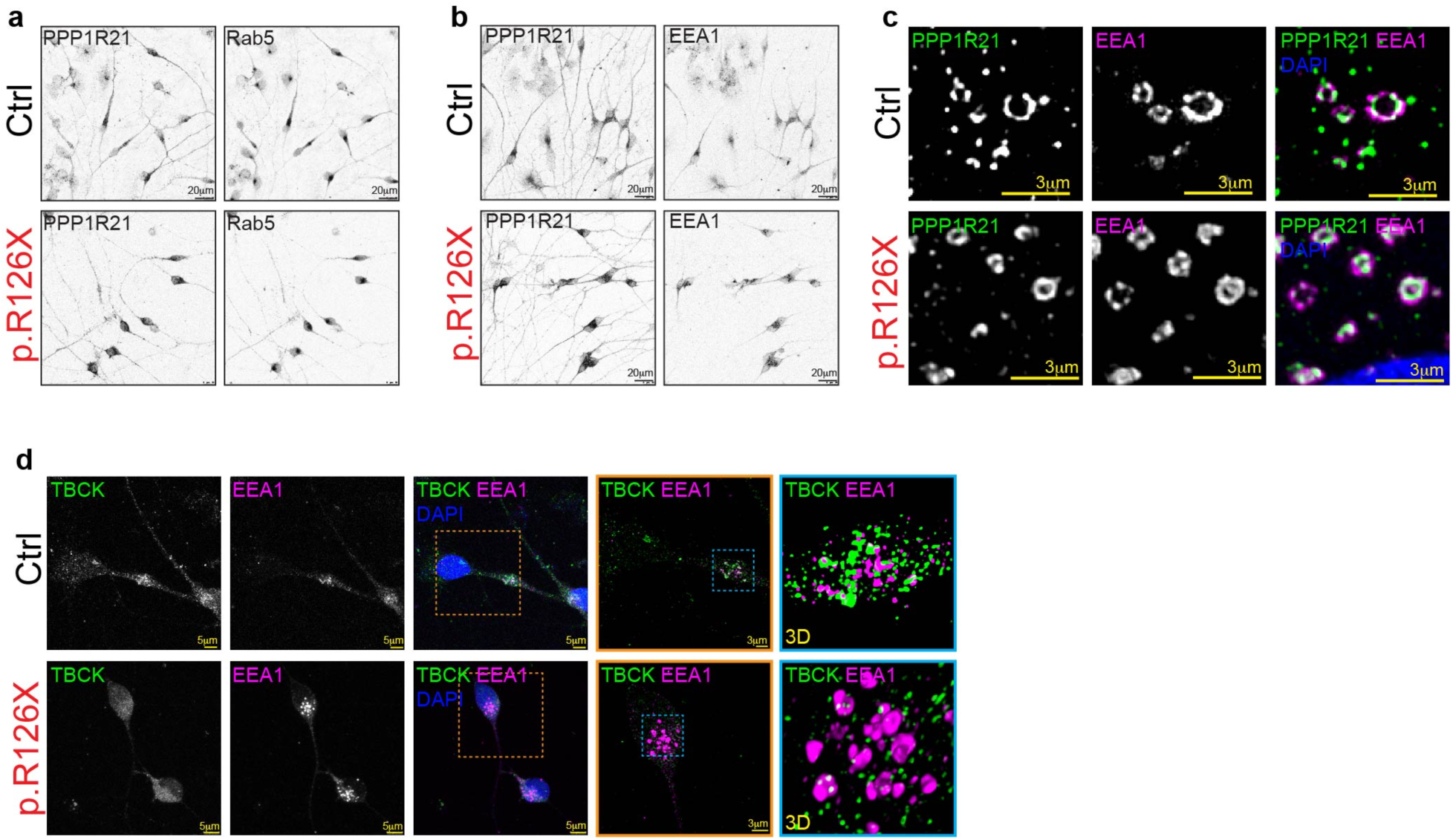
HEK cells deficient of TBCK show alterations in the distribution of PPP1R21 and early endosomes. **a,** Representative images showing the distribution of PPP1R21 with RAB5 or (**b**) EEA1 in control (Ctrl) TBCK-deficient neurons (p.R126X). **c,** HyVolution confocal images show the interaction between PPP1R21 and EEA1 in control (Ctrl) and TBCK-deficient fibroblasts (p.R126X). **d,** Confocal images show TBCK and EEA1 in control (Ctrl) TBCK-deficient neurons (p.R126X). Different inset magnifications (orange and blue squares) show the distribution and colocalization of TBCK and EEA1.

**Fig. S7:**
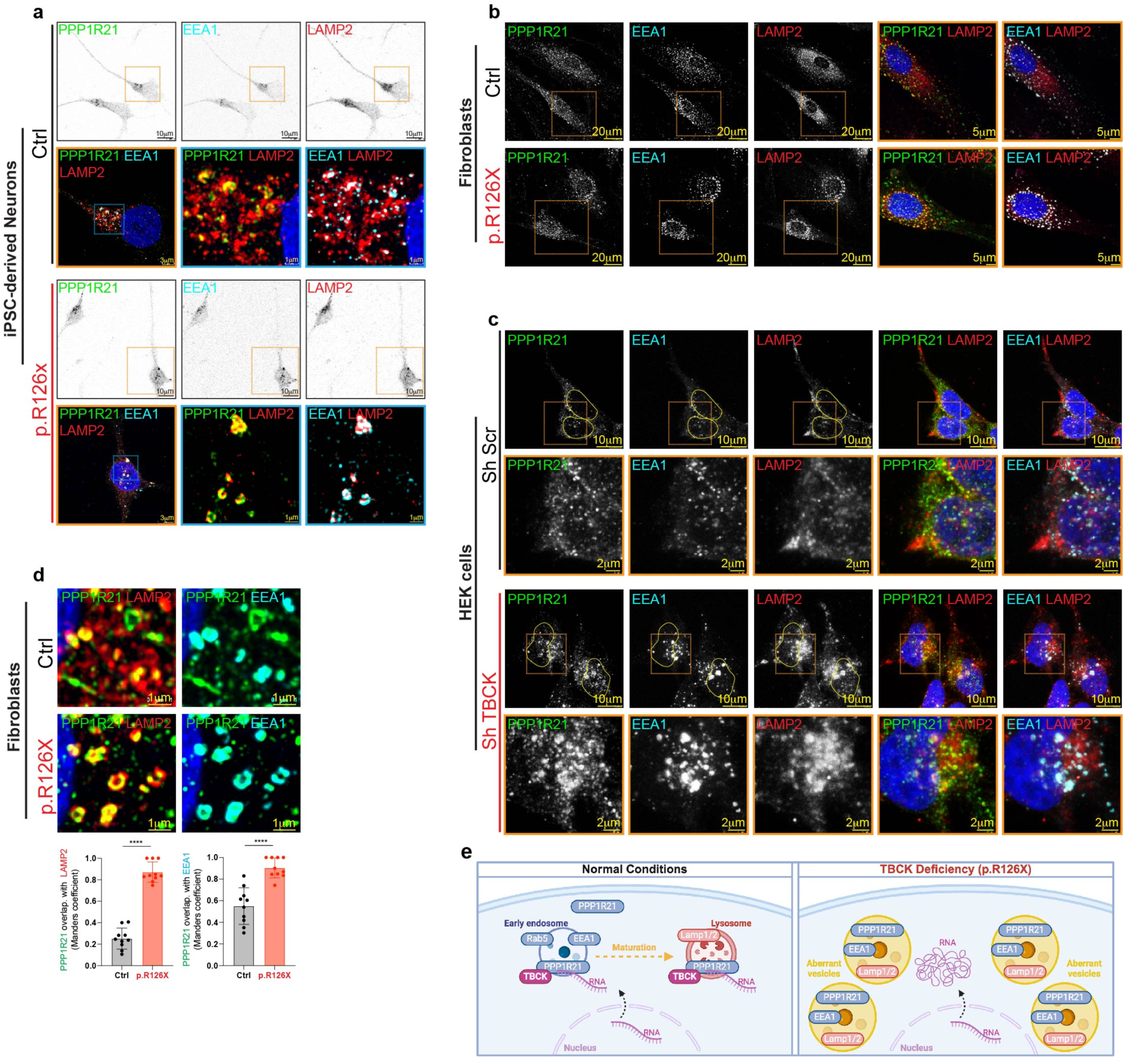
TBCK regulates interaction of PPP1R21 with endosomes and lysosomes. **a,** Representative confocal images in control (Ctrl) and TBCK-deficient neurons (p.R126X) showing PPP1R21, EEA1 (early endosomes) and LAMP2 (lysosomes). Inset magnifications (orange and blue squares) show the distribution and colocalization of PPP1R21, EEA1 and LAMP2. **b,** Selected confocal images show PPP1R21, EEA1 and LAMP2 in control (Ctrl) and TBCK-deficient fibroblasts (p.R126X). Different inset magnifications (orange squares) show the distribution and colocalization of PPP1R21, EEA1 and LAMP2. **c,** Confocal images of HEK293 cells transduced with ShRNA scramble (Sh Scr) and ShRNA TBCK#2 (Sh TBCK). Different inset magnifications (orange squares) show the distribution and colocalization of PPP1R21, EEA1 and LAMP2. **d,** Colocalization analysis using the Mander’s overlap coefficient (MOC) to quantity the degree of colocalization between PPP1R21 with LAMP2; LAMP2 with EEA1; and PPP1R21 with EEA1 in control and TBCK-deficient fibroblasts (p.R126X) (n=3). **e,** Function of TBCK in control (normal conditions) and TBCK-deficient cells. In presence of TBCK, one portion of PPP1R21 is localized in early endosomes with RAB5 and EEA1, but also another portion can be observed distributed in the cytoplasm. In absence of TBCK, PPP1R21 in mainly localized into large perinuclear vesicles formed by early endosomal (EEA1) and lysosomal (LAMP1/2) markers. This suggest that the PPP1R21 regulation by TBCK is essential for the formation, distribution or maturation of endosomes and lysosomes. These alterations impact also in the distribution and trafficking of mRNA mediated through the endolysosomal pathway which led to cellular defects and subsequently to TBCK Encephaloneuronopathy. p<0.01, **p<0.05 and ***p<0.001; Student’s t test; all graphs show error bars with mean ± SD from independent experiments (n).

## Notes

### Competing Interest Statement

The authors have declared no competing interest.

